# Cladogenetic and Anagenetic Models of Chromosome Number Evolution: a Bayesian Model Averaging Approach

**DOI:** 10.1101/086629

**Authors:** William A. Freyman, Sebastian Höhna

## Abstract

Chromosome number is a key feature of the higher-order organization of the genome, and changes in chromosome number play a fundamental role in evolution. Dysploid gains and losses in chromosome number, as well as polyploidization events, may drive reproductive isolation and lineage diversification. The recent development of probabilistic models of chromosome number evolution in the groundbreaking work by Mayrose et al. (2010, ChromEvol) have enabled the inference of ancestral chromosome numbers over molecular phylogenies and generated new interest in studying the role of chromosome changes in evolution. However, the ChromEvol approach assumes all changes occur anagenetically (along branches), and does not model events that are specifically cladogenetic. Cladogenetic changes may be expected if chromosome changes result in reproductive isolation. Here we present a new class of models of chromosome number evolution (called ChromoSSE) that incorporate both anagenetic and cladogenetic change. The ChromoSSE models allow us to determine the mode of chromosome number evolution; is chromosome evolution occurring primarily within lineages, primarily at lineage splitting, or in clade-specific combinations of both? Furthermore, we can estimate the location and timing of possible chromosome speciation events over the phylogeny. We implemented ChromoSSE in a Bayesian statistical framework, specifically in the software RevBayes, to accommodate uncertainty in parameter estimates while leveraging the full power of likelihood based methods. We tested ChromoSSE’s accuracy with simulations and re-examined chromosomal evolution in *Aristolochia, Carex* section *Spirostachyae, Helianthus, Mimulus* sensu lato (s.l.), and *Primula* section *Aleuritia*, finding evidence for clade-specific combinations of anagenetic and cladogenetic dysploid and polyploid modes of chromosome evolution.

---

A central organizing component of the higher-order architecture of the genome is chromosome number, and changes in chromosome number have long been understood to play a fundamental role in evolution. In the seminal work *Genetics and the Origin of Species* (1937), Dobzhansky identified “the raw materials for evolution”, the sources of natural variation, as two evolutionary processes: mutations and chromosome changes. “Chromosomal changes are one of the mainsprings of evolution,” Dobzhansky asserted, and changes in chromosome number such as the gain or loss of a single chromosome (dysploidy), or the doubling of the entire genome (polyploidy), can have phenotypic consequences, affect the rates of recombination, and increase reproductive isolation among lineages and thus drive diversification (Stebbins 1971). Recently, evolutionary biologists have studied the macroevolutionary consequences of chromosome changes within a molecular phylogenetic framework, mostly due to the groundbreaking work of Mayrose et al. (2010, ChromEvol) which introduced likelihood-based models of chromosome number evolution. The ChromEvol models have permitted phylogenetic studies of ancient whole genome duplication events, rapid “catastrophic” chromosome speciation, major reevaluations of the evolution of angiosperms, and new insights into the fate of polyploid lineages (e.g. Pires and Hertweck 2008; Mayrose et al. 2011; Tank et al. 2015).

One aspect of chromosome evolution that has not been thoroughly studied in a probabilistic framework is cladogenetic change in chromosome number. Cladogenetic changes occur solely at speciation events, as opposed to anagenetic changes that occur within lineages and are not associated with speciation events. Studying cladogenetic chromosome changes in a phylogenetic framework has been difficult since the approach used by ChromEvol models only anagenetic changes and ignores the changes that occur specifically at speciation events and may be expected if chromosome changes result in reproductive isolation. Reproductive incompatibilities caused by chromosome changes may play an important role in the speciation process, and led White (1978) to propose that chromosome changes perform “the primary role in the majority of speciation events.” Indeed, chromosome fusions and fissions may have played a role in the formation of reproductive isolation and speciation in the great apes (Ayala and Coluzzi 2005), and the importance of polyploidization in plant speciation has long been appreciated (Coyne et al. 2004; Rieseberg and Willis 2007). Recent work by Zhan et al. (2016) revealed phylogenetic evidence that polyploidization is frequently cladogenetic in land plants. However, their approach did not examine the role dysploid changes may play in speciation, and it required a two step analysis in which one first used ChromEvol to infer ploidy levels, and then a second modeling step to infer the proportion of ploidy shifts that were cladogenetic. Since ChromEvol only models anagenetic polyploidization events these two modeling steps are inconsistent with one another.

Here we present models of chromosome number evolution that simultaneously account for both cladogenetic and anagenetic polyploid as well as dysploid changes in chromosome number over a phylogeny. These models reconstruct an explicit history of cladogenetic and anagenetic changes in a clade, enabling estimation of ancestral chromosome numbers. Our approach also identifies different modes of chromosome number evolution among clades; we can detect primarily anagenetic, primarily cladogenetic, or clade-specific combinations of both modes of chromosome changes. Furthermore, these models allow us to infer the timing and location of possible polyploid and dysploid speciation events over the phylogeny. Since these models only account for changes in chromosome number, they ignore speciation that may accompany other types of chromosome rearrangements such as inversions. Our models cannot determine that changes in chromosome number “caused” the speciation event, but they do reveal that speciation and chromosome change are temporally correlated. Thus, these models can give us evidence that the chromosome number change coincided with cladogenesis and so may have played a significant role in the speciation process.

A major challenge for all phylogenetic models of cladogenetic character change is accounting for unobserved speciation events due to lineages going extinct and not leaving any extant descendants (Bokma 2002), or due to incomplete sampling of lineages in the present. Teasing apart the phylogenetic signal for cladogenetic and anagenetic processes given unobserved speciation events is a major difficulty. The Cladogenetic State change Speciation and Extinction (ClaSSE) model (Goldberg and Igić 2012) accounts for unobserved speciation events by jointly modeling both character evolution and the phylogenetic birth-death process. Our class of chromosome evolution models uses the ClaSSE approach, and could be considered a special case of ClaSSE. We implemented our models (called ChromoSSE) in a Bayesian framework and use Markov chain Monte Carlo algorithms to estimate posterior probabilities of the model’s parameters. However, compared to most character evolution models, SSE models require additional complexity since they must model extinction and speciation processes. Using simulations, we examined the impact of this additional complexity on our chromosome evolution models’ performance. Note that ChromoSSE uses the SSE approach to integrate over all unobserved speciation events and in this work we do not investigate how chromosome number affects diversification rates. Nonetheless, our implementation enables chromosome number dependent speciation and extinction rates to be estimated and this will be explored in future work.

Out of the class of ChromoSSE models described here, it is possible that no single model will adequately describe the chromosome evolution of a given clade. The most parameter-rich ChromoSSE model has at least 12 independent rate parameters, however the models that best describe a given dataset (a phylogeny and a set of observed chromosome counts) may be special cases of the full model. For example, there may be a clade for which the best fitting models have no anagenetic rate of polyploidization (the rate = 0.0) and for which all polyploidization events are cladogenetic. To explore the entire space of all possible models of chromosome number evolution we constructed a reversible jump Markov chain Monte Carlo (Green 1995) that samples across models of different dimensionality, drawing samples from chromosome evolution models in proportion to their posterior probability and enabling Bayes factors for each model to be calculated. This approach incorporates model uncertainty by permitting model-averaged inferences that do not condition on a single model; we draw estimates of ancestral chromosome numbers and rates of chromosome evolution from all possible models weighted by their posterior probability. For general reviews of this approach to model averaging see Madigan and Raftery (1994), Hoeting et al. (1999), Kass and Raftery (1995), and for its use in phylogenetics see Posada and Buckley (2004). Averaging over all models has been shown to provide a better average predictive ability than conditioning on a single model (Madigan and Raftery 1994). Conditioning on a single model ignores model uncertainty, which can lead to an underestimation in the uncertainty of inferences made from that model (Hoeting et al. 1999). In our case, this can lead to overconfidence in estimates of ancestral chromosome numbers and chromosome evolution parameter value estimates.

Our motivation in developing these phylogenetic models of chromosome evolution is to determine the mode of chromosome number evolution; is chromosome evolution occurring primarily within lineages, primarily at lineage splitting, or in clade-specific combinations of both? By identifying how much of the pattern of chromosome number evolution is explained by anagenetic versus cladogenetic change, and by identifying the timing and location of possible chromosome speciation events over the phylogeny, the ChromoSSE models can help uncover how much of a role chromosome changes play in speciation. In this paper we first describe the ChromoSSE models of chromosome evolution and our Bayesian method of model selection, then we assess the models’ efficacy by testing them with simulated datasets, particularly focusing on the impact of unobserved speciation events on inferences, and finally we apply the models to five empirical datasets that have been previously examined using other models of chromosome number evolution.

## Methods

### Models of Chromosome Evolution

In this section we introduce our class of probabilistic models of chromosome number evolution. We are interested in modeling the changes in chromosome number both within lineages (anagenetic evolution) and at speciation events (cladogenetic evolution). The anagenetic component of the model is a continuous-time Markov process similar to Mayrose et al. (2010) as described below. The cladogenetic changes are accounted for by a birth-death process similar to Maddison et al. (2007) and Goldberg and Igić (2012), except each type of cladogenetic chromosome event is given its own rate. Thus, the birth-death process has multiple speciation rates (one for each type of cladogenetic change) and a single constant extinction rate. Our models of chromosome number evolution can therefore be understood as a specific case of the Cladogenetic State change Speciation and Extinction (ClaSSE) model (Goldberg and Igić 2012), which integrates over all possible unobserved speciation events (due to lineages that were unsampled or have gone extinct) directly in the likelihood calculation of the observed chromosome counts and tree shape. To test the importance of accounting for unobserved speciation events we also briefly describe a version of the model that handles different cladogenetic event types as transition probabilities at each observed speciation event and ignores unobserved speciation events, similar to the dispersal-extinction-cladogenesis (DEC) models of geographic range evolution (Ree and Smith 2008).

Our implementation assumes chromosome numbers can take the value of any positive integer, however to limit the transition matrices to a reasonable size for likelihood calculations we follow Mayrose et al. (2010) in setting the maximum chromosome number *C_m_* to *n* + 10, where *n* is the highest chromosome number in the observed data. Note that we allow this parameter to be set in our implementation. Hence, it is easily possible to test the impact of setting a specific value for the maximum chromosome count.

Our models contain a set of 6 free parameters for anagenetic chromosome number evolution, a set of 5 free parameters for cladogenetic chromosome number evolution, an extinction rate parameter, and a vector of *C_m_* root frequencies of chromosome numbers, for a total of 12 + *C_m_* free parameters. All of the 11 chromosome rate parameters can be removed (fixed to 0.0) except the cladogenetic no-change rate parameter. Thus, the class of chromosome number evolution models described here has a total of 2^10^ = 1024 nested models of chromosome evolution.

#### Chromosome evolution within lineages

Chromosome number evolution within lineages (anagenetic change) is modeled as a continuous-time Markov process similar to Mayrose et al. (2010). The continuous-time Markov process is described by an instantaneous rate matrix *Q* where the value of each element represents the instantaneous rate of change within a lineage from a genome of *i* chromosomes to a genome of *j* chromosomes. For all elements of *Q* in which either *i* = 0 or *j* = 0 we define *Q*_*ij*_ = 0. For the off-diagonal elements *i* = *j* with positive values of *i* and *j, Q* is determined by:

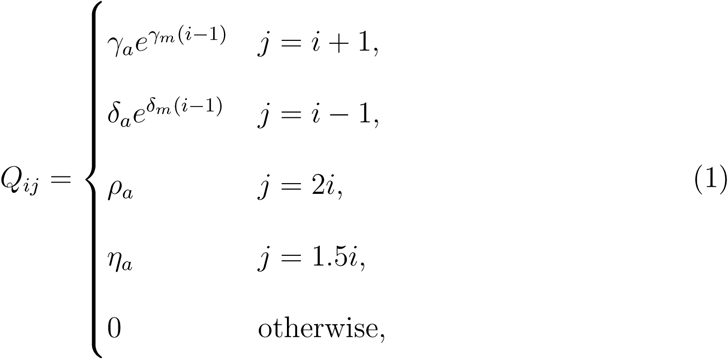

where *γ_a_*, *δ_a_*, *ρ_a_*, and *η_a_* are the rates of chromosome gains, losses, polyploidizations, and demi-polyploidizations. *γ_m_* and *δ_η_* are rate modifiers of chromosome gain and loss, respectively, that allow the rates of chromosome gain and loss to depend on the current number of chromosomes. This enables modeling scenarios in which the probability of fusion or fission events is positively or negatively correlated with the number of chromosomes. If the rate modifier *γ_m_* = 0, then *γ_a_e*^0(*i*−1)^ = *γ_a_*. If the rate modifier *γ_m_* > 0, then *γ_a_e*^*γ_m_*(*i*-1)^ ≥ *γ_a_*, and if *γ_m_* < 0 then *γ_a_e*^*γ_m_*(*i*−1)^ ≤ *γ_a_*. These two rate modifiers replace the parameters *λ_l_* and *δ_l_* in Mayrose et al. (2010), which in their parameterization may result in negative transition rates. Here we chose to exponentiate *γ_m_* and *δ_m_* to ensure positive transition rates, and avoid ad hoc restrictions on negative transition rates that may induce unintended priors. Note that this assumes the rates of chromosome change can vary exponentially as a function of the current chromosome number, whereas Mayrose et al. (2010) assumes a linear function.

For odd values of *i*, we set *Q*_*ij*_ = *η*/2 for the two integer values of *j* resulting when *j* = 1.5*i* was rounded up and down. We define the diagonal elements *i* = *j* of *Q* as: 
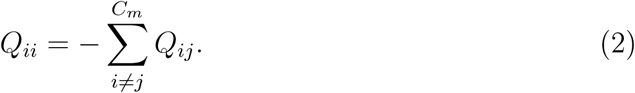

The probability of anagenetically transitioning from chromosome number *i* to *j* along a branch of length *t* is then calculated by exponentiation of the instantaneous rate matrix:

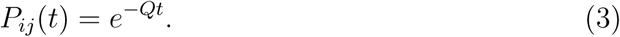

#### Chromosome evolution at cladogenesis events

At each lineage divergence event over the phylogeny, nine different cladogenetic changes in chromosome number are possible (Figure 1). Each type of cladogenetic event occurs with the rate *φ_c_,γ_c_,δ_c_, ρ_c_, η_c_*, representing the cladogenesis rates of no change, chromosome gain, chromosome loss, polyploidization, and demi-polyploidization, respectively. The speciation rates λ for the birth-death process generating the tree are given in the form of a 3-dimensional matrix between the ancestral state *i* and the states of the two daughter lineages *j* and *k*. For all positive values of *i, j*, and *k*, we define: 
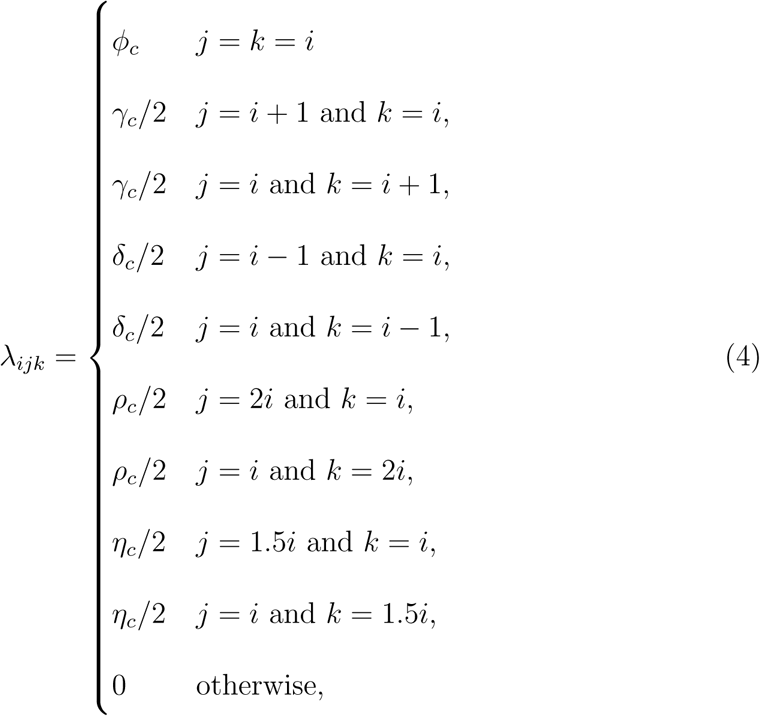
so that the total speciation rate of the birth-death process *λ_t_* is given by:

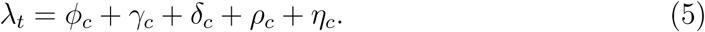

Similar to the anagenetic instantaneous rate matrix described above, for odd values of *i*, we set *λ*_*ijk*_ = *η_c_*/4 for the integer values of *j* and *k* resulting when 1.5*i* is rounded up and down. The extinction rate *μ* is constant over the tree and for all chromosome numbers.

**Figure 1:**
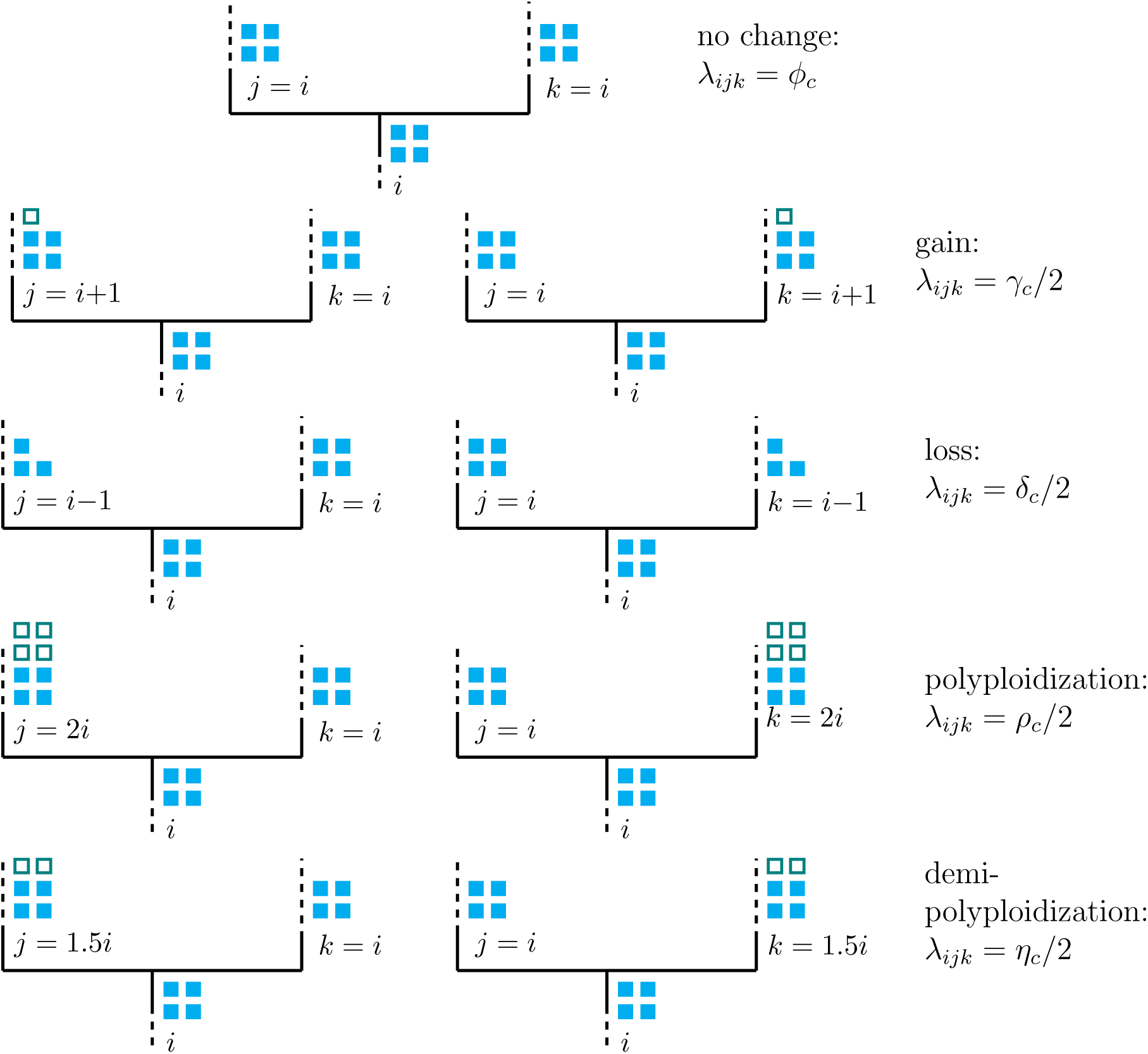
Modeled cladogenetic chromosome evolution events. At each speciation event 9 different cladogenetic events are possible. The rate of each type of speciation event is *λ_ijk_* where *i* is the chromosome number before cladogenesis and *j* and *k* are the states of each daughter lineage immediately after cladogenesis. The dashed lines represent possible chromosomal changes within lineages that are modeled by the anagenetic rate matrix *Q*.

Note that this model allows only a single chromosome number change event on a maximum of one of the daughter lineages at each cladogenesis event. Changes in both daughter lineages at cladogenesis are not allowed; at least one of the daughter lineages must inherit the chromosome number of the ancestor. The model also assumes that cladogenesis events are always strictly bifurcating and that there are no hard polytomies.

#### Likelihood Calculation Accounting for Unobserved Speciation

The likelihood of cladogenetic and anagenetic chromosome number evolution over a phylogeny is calculated using a set of ordinary differential equations similar to the Binary State Speciation and Extinction (BiSSE) model (Maddison et al. 2007). The BiSSE model was extended to incorporate cladogenetic changes by Goldberg and Igić (2012). Following Goldberg and Igić (2012), we define *D_Ni_*(*t*) as the probability that a lineage with chromosome number *i* at time *t* evolves into the observed clade *N*. We let *E_i_*(*t*) be the probability that a lineage with chromosome number *i* at time *t* goes extinct before the present, or is not sampled at the present. However, unlike the full ClaSSE model the extinction rate *μ* does not depend on the chromosome number *i* of the lineage. The differential equations for these two probabilities is given by: 
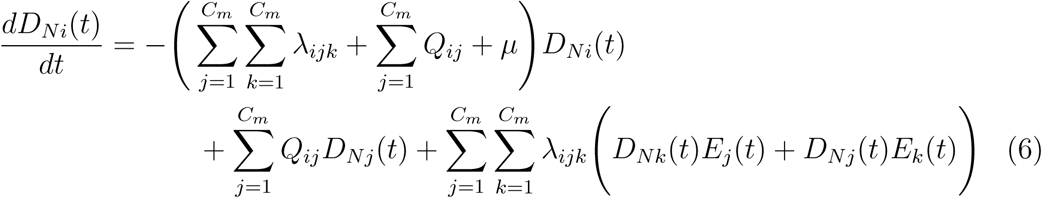

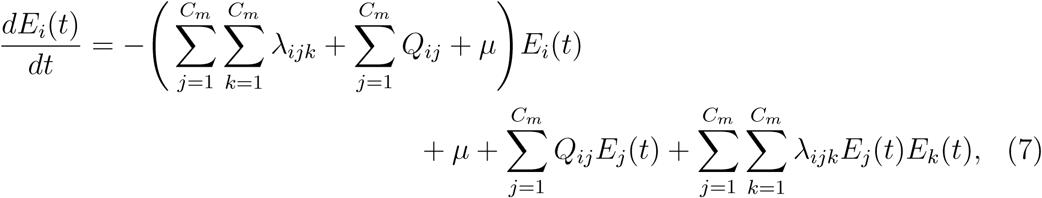
where λ_*ijk*_ for each possible cladogenetic event is given by equation 4, and the rates of anagenetic changes *Q*_*ij*_ are given by equation 1. See Figure 2 for an explanation of equations 6 and 7.

**Figure 2:**
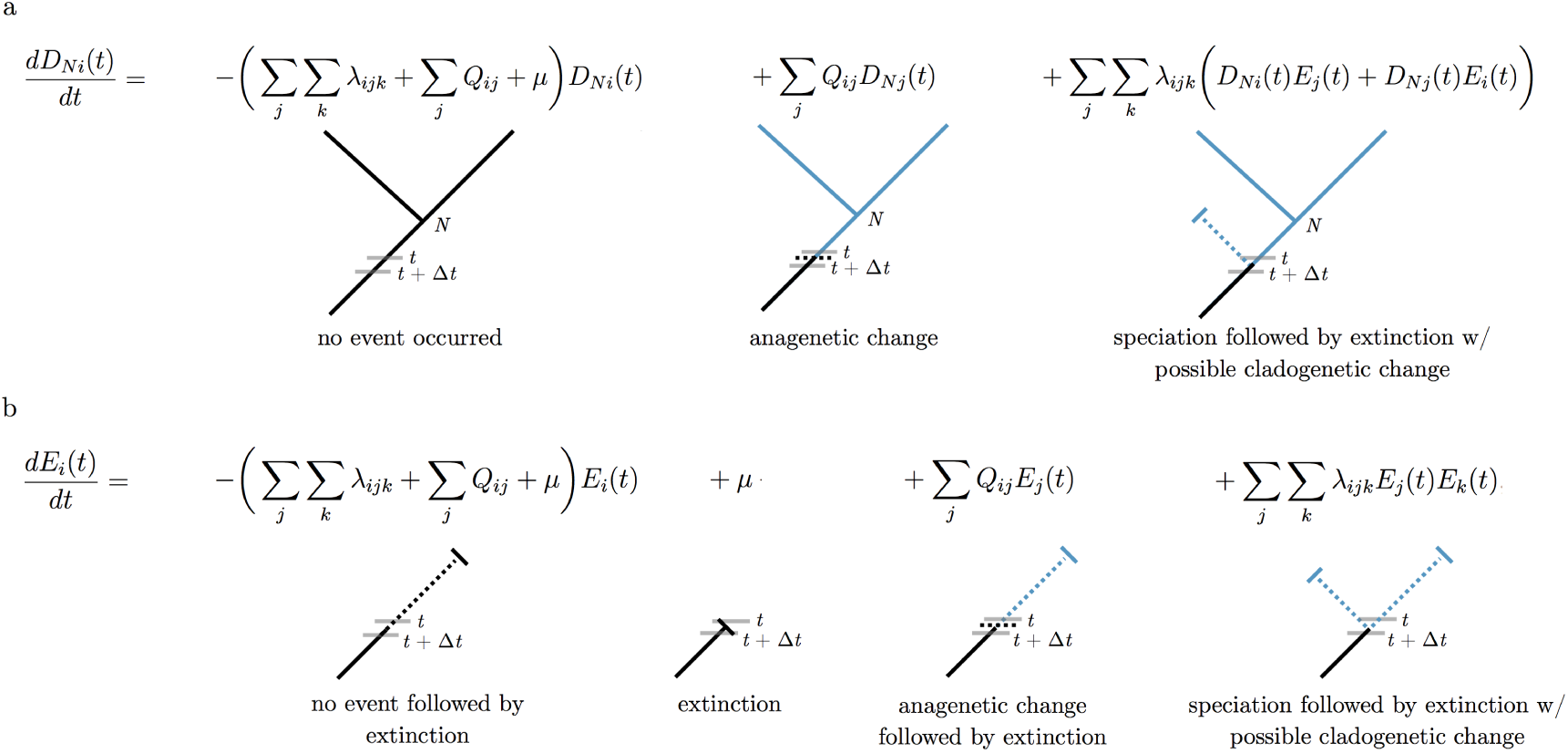
Chromosome evolution through time. An illustration of chromosome evolution events that could occur during each time interval *Δt* along the branches of a phylogeny. Equations 6 and 7 (subfigures a and b, respectively) sum over each possible chromosome evolution event and are numerically integrated backwards through time over the phylogeny to calculate the likelihood. a) *D_Ni_*(*t*) is the probability that the lineage at time *t* evolves into the observed clade *N*. To calculate the change in this probability over Δ*t* we sum over three possibilities: no event occurred, an anagenetic change in chromosome number occurred, or a speciation event with a possible cladogenetic chromosome change occurred followed by an extinction event on one of the two daughter lineages. b) *E_i_*(*t*) is the probability that the lineage goes extinct or is not sampled at the present. To calculate the change in this probability over Δ*t* we sum over four possibilities: no event occurred followed eventually by extinction, extinction occurred, an anagenetic change occurred followed by extinction, or a speciation event with a possible cladogenetic change occurred followed by extinction of both daughter lineages.

The differential equations above have no known analytical solution. Therefore, we numerically integrate the equations for every arbitrarily small time interval moving along each branch from the tip of the tree towards the root. When a node *l* is reached, the probability of it being in state *i* is calculated by combining the probabilities of its descendant nodes *m* and *n* as such: 
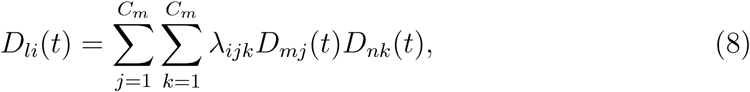
where again λ_*ijk*_ for each possible cladogenetic event is given by equation 4. Letting *D* denote a set of observed chromosome counts, Ψ an observed phylogeny, and *θ_q_* a particular set of chromosome evolution model parameters, then the likelihood for the model parameters *θ_q_* is given by: 
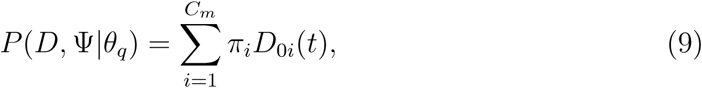
where *π_i_* is the root frequency of chromosome number *i* and *D_0i_*(*t*) is the likelihood of the root node being in state *i* conditional on having given rise to the observed tree Ψ and the observed chromosome counts *D*.

#### Initial Conditions

The initial conditions for each observed lineage at time *t* = 0 for the extinction probabilities described by equation 7 are *E_i_*(0) = 1 − *ρ_s_* for all *i* where *ρ_s_* is the sampling probability of including that lineage. For lineages with an observed chromosome number of *i*, the initial condition is *D_Ni_*(0) = *ρ_s_*. The initial condition for all other chromosome numbers *j* is *D_Nj_*(0) = 0.

#### Likelihood Calculation Ignoring Unobserved Speciation

To test the effect of unobserved speciation events on inferences of chromosome number evolution we also implemented a version of the model described above that only accounts for observed speciation events. At each lineage divergence event over the phylogeny, the probabilities of cladogenetic chromosome number evolution *P*({*j,k*}|*i*) are given by the simplex {*φ_ρ_, γ_ρ_, δ_ρ_, p_p_, η_ρ_*}, where *φ_ρ_, γ_ρ_, δ_ρ_, p_p_*, and *η_ρ_* represent the probabilities of no change, chromosome gain, chromosome loss, polyploidization, and demi-polyploidization, respectively. This approach does not require estimating speciation or extinction rates.

Here, we calculate the likelihood of chromosome number evolution over a phylogeny using Felsenstein’s pruning algorithm (Felsenstein 1981) modified to include cladogenetic probabilities similar to models of biogeographic range evolution (Landis et al. 2013; Landis in press). Let *D* again denote a set of observed chromosome counts and Ψ represent an observed phylogeny where node *l* has descendant nodes *m* and *n*. The likelihood of chromosome number evolution at node *l* conditional on node *l* being in state *i* and *θ_q_* being a particular set of chromosome evolution model parameter values is given by: 
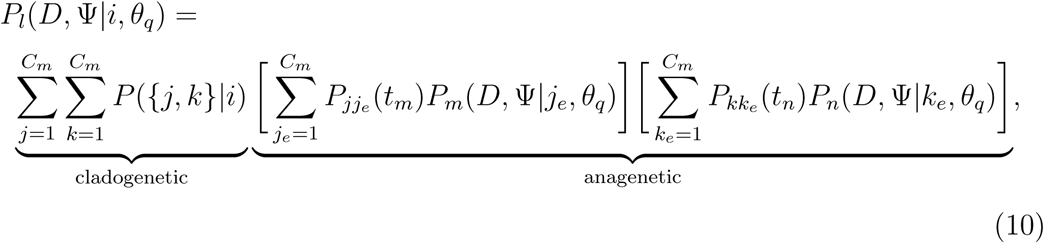
where the length of the branches between *l* and *m* is *t_m_* and between *l* and *n* is *t_n_*. The state at the end of these branches near nodes *m* and *n* is *j_e_* and *k_e_*, respectively. The state at the beginning of these branches, where they meet at node *l*, is *j* and *k* respectively. The cladogenetic term sums over the probabilities *P*({*j, k*}|*i*) of all possible cladogenetic changes from state *i* to the states *j* and *k* at the beginning of each daughter lineage. The anagenetic term of the equation is the product of the probability of changes along the branches from state *j* to state *j_e_* and state *k* to state *k_e_* (given by equation 3) and the likelihood of the tree above node *l* recursively computed from the tips.

The likelihood for the model parameters *θ_q_* is given by: 
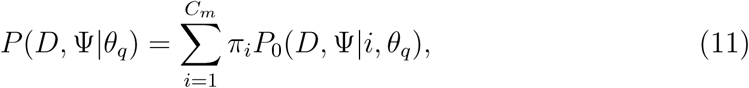
where *P*_0_(*D*, Ψ| *i*, *θ_q_*) is the conditional likelihood of the root node being in state *i* and *π_i_* is the root frequency of chromosome number *i*.

#### Estimating Parameter Values and Ancestral States

For any given tree with a set of observed chromosome counts, there exists a posterior distribution of model parameter values and a set of probabilities for the ancestral chromosome numbers at each internal node of the tree. Let *P*(*s_i_, θ_q_* |*D*, Ψ) denote the joint posterior probability of *θ_q_* and a vector of specific ancestral chromosome numbers *s_i_* given a set of observed chromosome counts *D* and an observed tree Ψ. The posterior is given by Bayes’ rule:

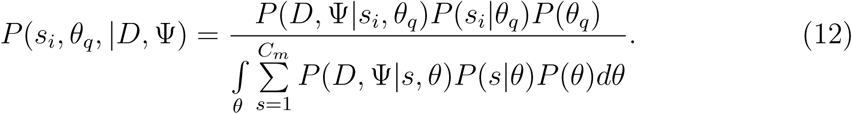

Here, *P*(*s_i_|θ_q_*) is the prior probability of the ancestral states *s* conditioned on the model parameters *θ_q_*, and *P*(*θ_q_*) is the joint prior probability of the model parameters.

In the denominator of equation 12 we integrate over all possible values of *θ* and sum over all possible ancestral chromosome numbers *s*. Since *θ* is a vector of 12 + *C_m_* parameters and *s* is a vector of *n –* 1 ancestral states where *n* is the number of observed tips in the phylogeny, the denominator of equation 12 requires a high dimensional integral and an extremely large summation that is impossible to calculate analytically. Instead we use Markov chain Monte Carlo methods (Metropolis et al. 1953; Hastings 1970) to estimate the posterior probability distribution in a computationally efficient manner.

Ancestral states are inferred using a two-pass tree traversal procedure as described in Pupko et al. (2000), and previously implemented in a Bayesian framework by Huelsenbeck and Bollback (2001) and Pagel et al. (2004). First, partial likelihoods are calculated during the backwards-time post-order tree traversal in equations 6 and 7. Joint ancestral states are then sampled during a pre-order tree traversal in which the root state is first drawn from the marginal likelihoods at the root, and then states are drawn for each descendant node conditioned on the state at the parent node until the tips are reached. Again, we must numerically integrate over a system of differential equations during this root-to-tip tree traversal. This integration, however, is performed in forward-time, thus the set of ordinary differential equations must be slightly altered since our models of chromosome number evolution are not time reversible. Accordingly, we calculate: 
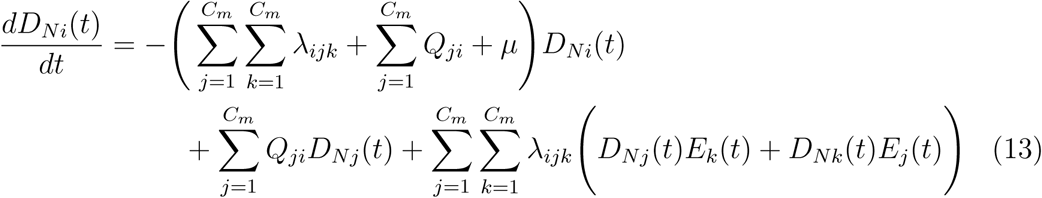

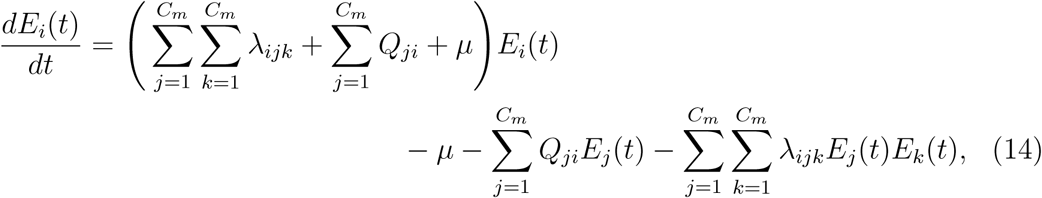
during the forward-time root-to-tip pass to draw ancestral states from their joint distribution conditioned on the model parameters and observed chromosome counts. For more details and validation of our method to estimate ancestral states, please see Supplementary Material Appendix 1.

#### Priors

Model parameter priors are listed in Table 1. Our implementation allows all priors to be easily modified so that their impact on results can be e**ff**ectively assessed. Priors for anagenetic rate parameters are given an exponential distribution with a mean of 2/Ψ_*l*_ where Ψ_*l*_ is the length of the tree Ψ. This corresponds to a mean rate of two events over the observed tree. The priors for the rate modifiers *γ_m_* and *δ_m_* are assigned a uniform distribution with the range –3/*C_M_* to 3/*C_m_*. This sets minimum and maximum bounds on the amount the rate modifiers can a**ff**ect the rates of gain and loss at the maximum chromosome number to *γ_a_e*^-3^ = *γ_a_*0.050 and *γ_a_e*^3^ = *γ_a_*20.1, and *δ_a_e*^-3^ = *δ_a_*0.050 and *δ_a_e*^3^ = *δ_a_*20.1, respectively.

**Table 1:**
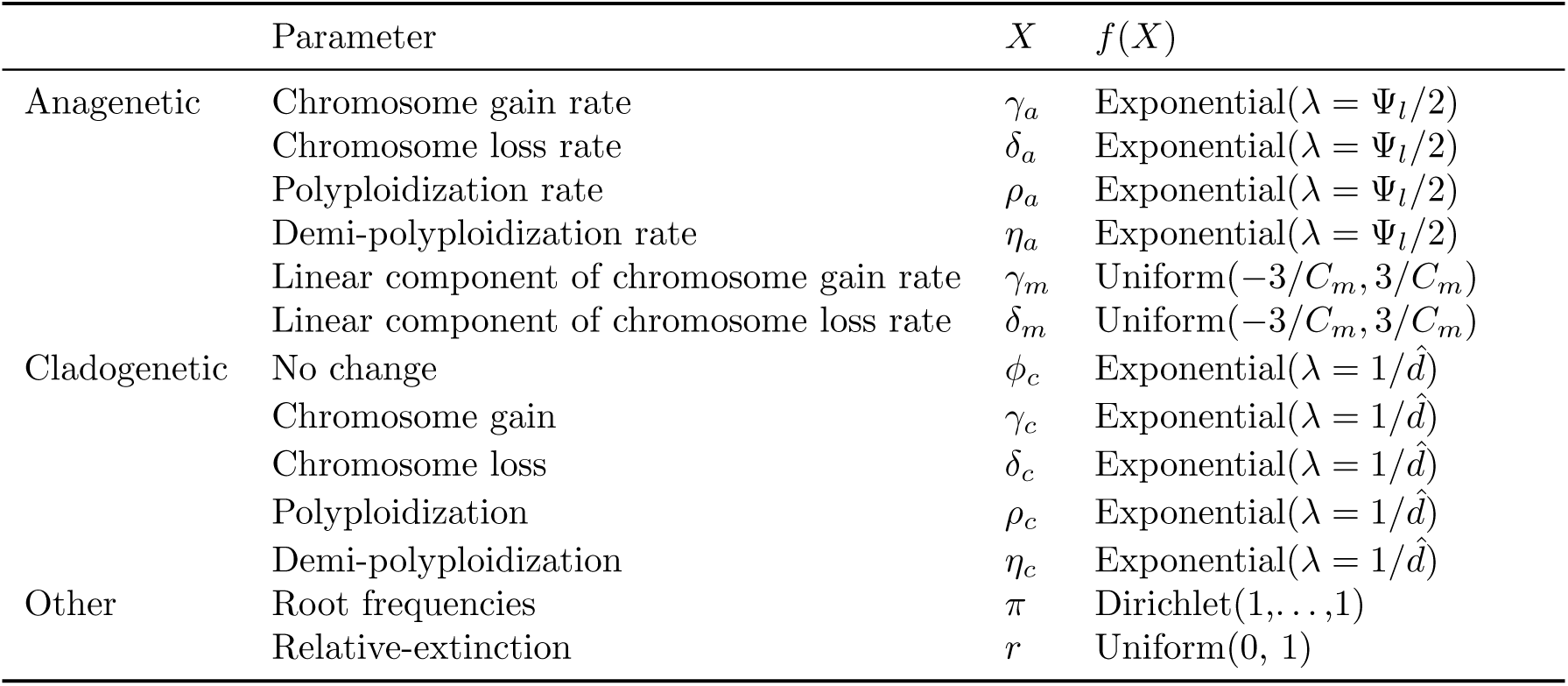
Model parameter names and prior distributions. See the main text for complete description of model parameters and prior distributions. Ψ_*l*_ represents the length of tree Ψ and *C_m_* is the maximum chromosome number allowed.

| | Parameter | $X$ | $f(X)$ |
| --- | --- | --- | --- |
| Anagenetic | Chromosome gain rate | $\gamma_a$ | Exponential($\lambda = \Psi_l/2$) |
| | Chromosome loss rate | $\delta_a$ | Exponential($\lambda = \Psi_l/2$) |
| | Polyploidization rate | $\rho_a$ | Exponential($\lambda = \Psi_l/2$) |
| | Demi-polyploidization rate | $\eta_a$ | Exponential($\lambda = \Psi_l/2$) |
| | Linear component of chromosome gain rate | $\gamma_m$ | Uniform($-3/C_m, 3/C_m$) |
| | Linear component of chromosome loss rate | $\delta_m$ | Uniform($-3/C_m, 3/C_m$) |
| Cladogenetic | No change | $\phi_c$ | Exponential($\lambda = 1/\hat{d}$) |
| | Chromosome gain | $\gamma_c$ | Exponential($\lambda = 1/\hat{d}$) |
| | Chromosome loss | $\delta_c$ | Exponential($\lambda = 1/\hat{d}$) |
| | Polyploidization | $\rho_c$ | Exponential($\lambda = 1/\hat{d}$) |
| | Demi-polyploidization | $\eta_c$ | Exponential($\lambda = 1/\hat{d}$) |
| Other | Root frequencies | $\pi$ | Dirichlet(1, ..., 1) |
| | Relative-extinction | $r$ | Uniform(0, 1) |

The speciation rates are drawn from an exponential prior with a mean equal to an estimate of the net diversification rate 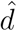. Under a constant rate birth-death process not conditioning on survival of the process, the expected number of lineages at time *t* is given by: 
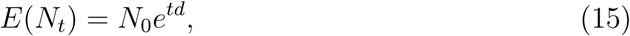
where *N_0_* is the number of lineages at time 0 and *d* is the net diversification rate *λ − μ* (Nee et al. 1994; Höhna 2015). Therefore, we estimate *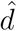* as:

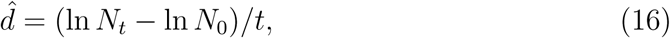
where *N_t_* is the number of lineages in the observed tree that survived to the present, *t* is the age of the root, and *N_0_* = 2.

The extinction rate *μ* is given by:

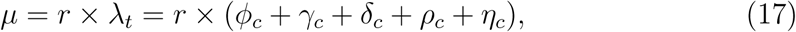

where *λ_t_* is the total speciation rate and *r* is the relative extinction rate. The relative extinction rate *r* is assigned a uniform (0,1) prior distribution, thus forcing the extinction rate to be smaller than the total speciation rate. The root frequencies of chromosome numbers π are drawn from a flat Dirichlet distribution.

### Model Uncertainty and Selection

#### Model Averaging

To account for model uncertainty we calculate the posterior density of chromosome evolution model parameters *θ* without conditioning on any single model of chromosome evolution. For each of the 1024 chromosome models *M_k_*, where *k* = 1, 2,…, 1024, the posterior distribution of *θ* is 
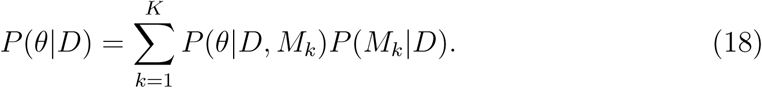

Here we average over the posterior distributions conditioned on each model weighted by the model’s posterior probability. We assume an equal prior probability for each model *P*(*M_k_*) = 2^-10^.

#### Reversible Jump Markov Chain Monte Carlo

To sample from the space of all possible chromosome evolution models, we employ reversible jump MCMC (Green 1995). This algorithm draws samples from parameter spaces of differing dimensions, and in stationarity samples each model in proportion to its posterior probability. This permits inference of each model’s fit to the data while simultaneously accounting for model uncertainty.

Our reversible jump MCMC moves between models of different dimensions using augment and reduce moves (Huelsenbeck et al. 2000; Pagel and Meade 2006; May et al. 2016). The reduce move proposes that a parameter should be removed from the current model by setting its value to 0.0, effectively disallowing that class of evolutionary event. Augment moves reverse reduce moves by allowing the parameter to once again have a non-zero value. Both augment and reduce moves operate on all chromosome rate parameters except for *φ* the rate of no cladogenetic change. Thus the least complex model the MCMC can sample from is one in which *φ_c_* > 0.0 and all other chromosome rate parameters are set to 0.0, corresponding to a model of no chromosomal changes over the phylogeny. The prior probability of reducing or augmenting model *M_k_* is *P_r_*(*M_k_*) = *P_a_*(*M_k_*) = 0.5.

#### Bayes Factors

In some cases we wish to compare the fit of models to summarize the mode of evolution within a clade. Bayes factors (Kass and Raftery 1995) compare the evidence between two competing models *M_i_* and *M_j_*

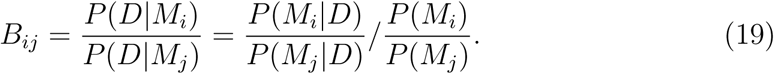

In words, the Bayes factor *B_ij_* is given by the ratio of the posterior odds to the prior odds of the two models. Unlike other methods of model selection such as Akaike Information Criterion (AIC; Akaike 1974) and the Bayesian Information Criterion (BIC; Schwarz 1978), Bayes factors take into account the full posterior densities of the model parameters and do not rely on point estimates. Furthermore AIC and BIC ignore the priors assigned to parameters, whereas Bayes factors penalizes parameters based on the informativeness of the prior. If the prior is informative but overlaps little with the likelihood it is penalized more than a diffuse uninformative prior that allows the parameter to take on whatever value is informed by the data (Xie et al. 2011).

### Implementation

The model and MCMC analyses described here are implemented in C++ in the software RevBayes (Höhna et al. 2016). In Supplementary Material Appendix 1 we validated our SSE likelihood calculations and ancestral state estimates against those of the R package diversitree (FitzJohn 2012). Rev scripts that specify the chromosome number evolution model (ChromoSSE) described here as a probabilistic graphical model (Höhna et al. 2014) and run the empirical analyses in RevBayes are available at http://github.com/wf8/ChromoSSE. The RevGadgets R package (available at https://github.com/revbayes/RevGadgets) contains functions to summarize results and generate plots of inferred ancestral chromosome numbers over a phylogeny.

The MCMC proposals used are outlined in Supplementary Material Appendix 2. Aside from the reversible jump MCMC proposals described above, all other proposals are standard except for the ElementSwapSimplex move operated on the Dirichlet distributed root frequencies parameter. This move randomly selects two elements *r*_1_ and *r*_2_ from the root frequencies vector and swaps their values. The reverse move, swapping the original values of *r*_1_ and *r*_2_ back, will have the same probability as the initial move since *r*_1_ and *r*_2_ were drawn from a uniform distribution. Thus, the Hasting ratio is 1 and the ElementSwapSimplex move is a symmetric Metropolis move.

### Simulations

We conducted a series of simulations to: 1) test the effect of unobserved speciation events due to extinction on chromosome number estimates when using a model that does not account for unobserved speciation, 2) compare the accuracy of models of chromosome evolution that account for unobserved speciation versus those that do not, 3) test the effect of jointly estimating speciation and extinction rates with chromosome number evolution, 4) test for identifiability of cladogenetic parameters, and 5) test the effect of incomplete sampling of extant lineages on ancestral chromosome number estimates. We will refer to each of the 5 simulations above as experiment 1, experiment 2, experiment 3, experiment 4, and experiment 5. Detailed descriptions of each experiment and the methods used to simulate trees and chromosome counts are in Supplementary Material Appendix 3.

For all 5 experiments, MCMC analyses were run for 5000 iterations, where each iteration consisted of 28 different moves in a random move schedule with 79 moves per iteration (see Supplementary material Appendix 2). Samples were drawn with each iteration, and the first 1000 samples were discarded as burn in. Effective sample sizes (ESS) for all parameters in all simulation replicates were over 200, and the mean ESS values of the posterior for the replicates was 1470.3. See Supplementary Material Appendix 4 for more on convergence of simulation replicates. To perform all 5 experiments 2100 independent MCMC analyses were run requiring a total of 89170.6 CPU hours on the Savio computational cluster at the University of California, Berkeley.

#### Summarizing Simulation Results

To summarize the results of our simulations, we measured the accuracy of ancestral state estimates as the percent of simulation replicates in which the true root chromosome number 8 was found to be the maximum a posteriori (MAP) estimate. To evaluate the uncertainty of the simulations, we calculated the mean posterior probability of root chromosome number for the simulation replicates that correctly found 8 to be the MAP estimate. We also calculated the proportion of simulation replicates for which the true model of chromosome number evolution used to simulate the data (as given by the table in Supplementary Material Appendix 3) was estimated to be the MAP model, and calculated the mean posterior probabilities of the true model. To compare the accuracy of model averaged parameter value estimates we calculated coverage probabilities. Coverage probabilities are the percentage of simulation replicates for which the true parameter value falls within the 95% highest posterior density (HPD). High accuracy is shown when coverage probabilities approach 1.0.

### Empirical Data

Phylogenetic data and chromosomes counts from five plant genera were analyzed (see Table 2). Like in Mayrose et al. (2010) we assumed each species had a single cytotype, however polymorphism could be accounted for by a vector of probabilities for each chromosome count. Sequence data for *Aristolochia* was downloaded from TreeBASE (Vos et al. 2010) study ID 1586. Sequences for *Helianthus, Mimulus* sensu lato, and *Primula* were downloaded directly from GenBank (Benson et al. 2005), reconstructing the sequence matrices from Timme et al. (2007), Beardsley et al. (2004), and Guggisberg et al. (2009). For each of these four datasets phylogenetic analyses were performed with all gene regions concatenated and unpartitioned, assuming the general time-reversible (GTR) nucleotide substitution model (Tavaré 1986; Rodriguez et al. 1990) with among-site rate variation modeled using a discretized gamma distribution (Yang 1994) with four rate categories. Since divergence time estimation in years is not the objective of this study, and only relative branching times are needed for our models of chromosome number evolution, a birth-death tree prior was used with a fixed root age of 10.0 time units. The MCMC analyses were performed in RevBayes, and were sampled every 100 iterations and run for a total of 400000 iterations, with samples from the first 100000 iterations discarded as burnin. Convergence was assessed by ensuring that the effective sample size for all parameters was over 200. The maximum a posteriori tree was calculated and used for further chromosome evolution analyses. For *Carex* section *Spirostachyae* the time calibrated tree from Escudero et al. (2010) was used.

**Table 2:**
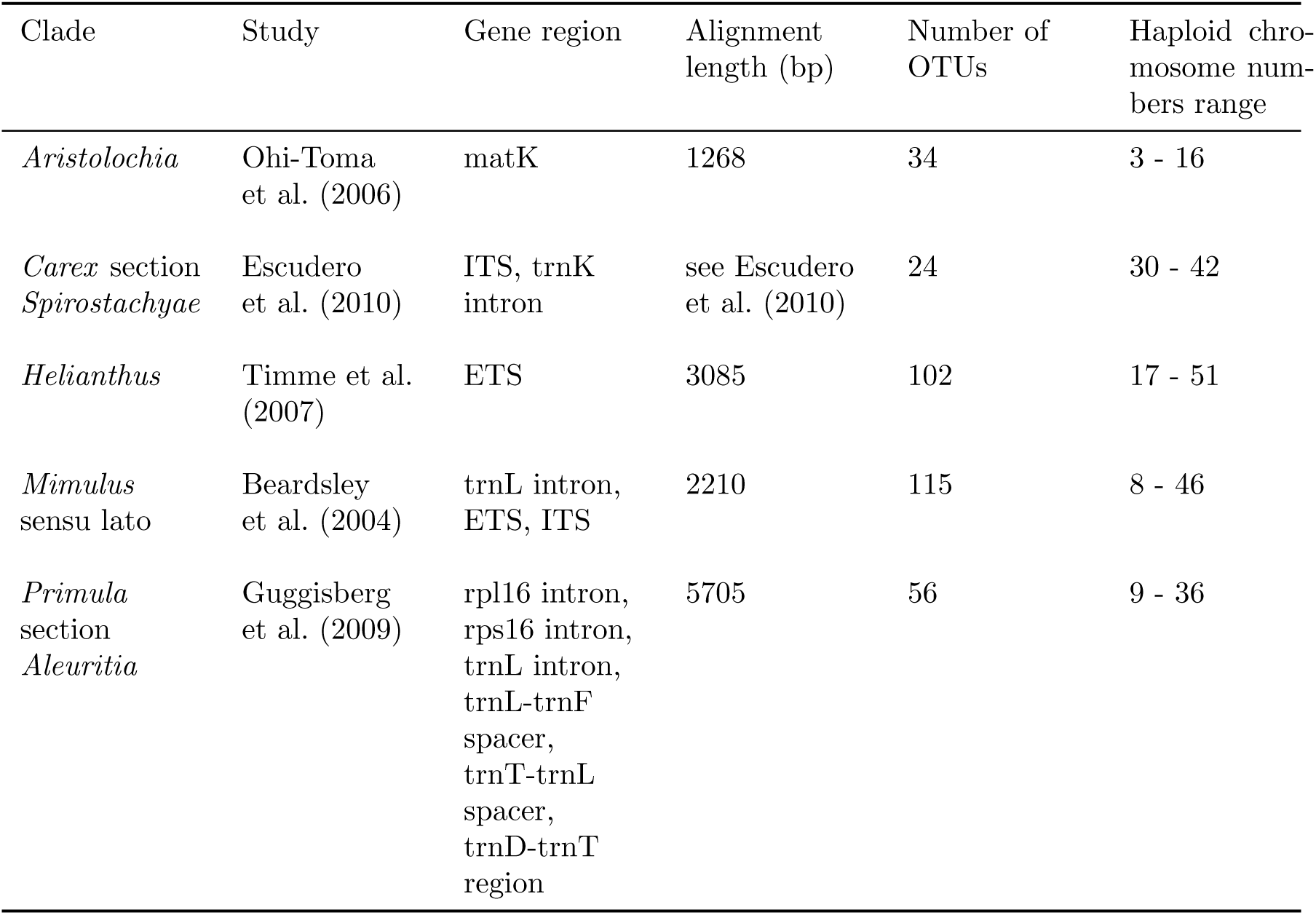
Empirical data sets analysed.

Ancestral chromosome numbers and chromosome evolution model parameters were then estimated for each of the five clades. Since testing the effect of incomplete taxon sampling on chromosome evolution inference of the empirical datasets was not a goal of this work, we focus here on results using a taxon sampling fraction *ρ_s_* of 1.0 (though see the Discussion section for more on this). MCMC analyses were run in RevBayes for 11000 iterations, where each iteration consisted of 28 different Metropolis-Hastings moves in a random move schedule with 79 moves per iteration (see Supplementary Material Appendix 2). Samples were drawn each iteration, and the first 1000 samples were discarded as burn in. Effective sample sizes for all parameters were over 200. For all datasets except *Primula* we used priors as outlined in Table 1. To demonstrate the flexibility of our Bayesian implementation and its capacity to incorporate prior information we used an informative prior for the root chromosome number in the *Primula* section *Aleuritia* analysis. Our dataset for *Primula* section *Aleuritia* also included samples from *Primula* sections *Armerina* and *Sikkimensis*. Since we were most interested in estimating chromosome evolution within section *Aleuritia*, we used an informative Dirichlet prior {1,…, 1,100, 1….1} (with 100 on the 11th element) to bias the root state towards the reported base number of *Primula x* =11 (Conti et al. 2000). Note all priors can be easily modified in our implementation, thus the impact of priors can be efficiently tested.

## Results

### Simulations

#### General Results

In all simulations, the true model of chromosome number evolution was infrequently estimated to be the MAP model (< 36% of replicates), and when it was the posterior probability of the MAP model was very low (< 0.12; Table 3). We found that the accuracy of root chromosome number estimation was similar whether the process that generated the simulated data was cladogenetic-only or anagenetic-only (Tables 3 and 4). However, when the data was simulated under a process that included both cladogenetic and anagenetic evolution we found a decrease in accuracy in the root chromosome number estimates in all cases.

**Table 3:** Experiment 1 results: the effect of ignoring unobserved speciation events on chromosome evolution estimates. Regardless of the true mode of chromosome evolution, the presence of unobserved speciation events in the process that generated the simulated data decreased accuracy in estimating the true root state. The columns from left to right are: 1) an indication of whether or not the data was simulated with a process that included unobserved speciation, 2) the true mode of chromosome evolution used to simulate the data, (for description see main text and Supplementary Material Appendix 3), 3) the percent of simulation replicates in which the true chromosome number at the root used to simulate the data was found to be the maximum a posteriori (MAP) estimate, 4) the mean posterior probability of the MAP estimate of the true root chromosome number, 5) the percent of simulation replicates in which the true model used to simulate the data was also found to be the MAP model, and 6) the mean posterior probability of the MAP estimate of the true model.

| Unobserved Speciation Events Included When Simulating Data? | Mode of Evolution Used to Simulate Data | True Root State Estimated (%) | Mean Posterior of True Root State | True Model Estimated (%) | Mean Posterior of True Model |
| --- | --- | --- | --- | --- | --- |
| No | Cladogenetic | 93 | 0.92 | 13 | 0.10 |
| No | Anagenetic | 89 | 0.91 | 31 | 0.12 |
| No | Mixed | 88 | 0.84 | 0 | 0.0 |
| Yes | Cladogenetic | 78 | 0.87 | 15 | 0.09 |
| Yes | Anagenetic | 83 | 0.91 | 36 | 0.12 |
| Yes | Mixed | 62 | 0.80 | 2 | 0.10 |

#### Experiment 1 Results

The presence of unobserved speciation in the process that generated the simulated data decreased the accuracy of ancestral state estimates (Figure 3, Table 3). Similarly, uncertainty in root chromosome number estimates increased with unobserved speciation (lower mean posterior probabilities; Table 3). The accuracy of parameter value estimates as measured by coverage probabilities was similar (results not shown).

**Figure 3:**
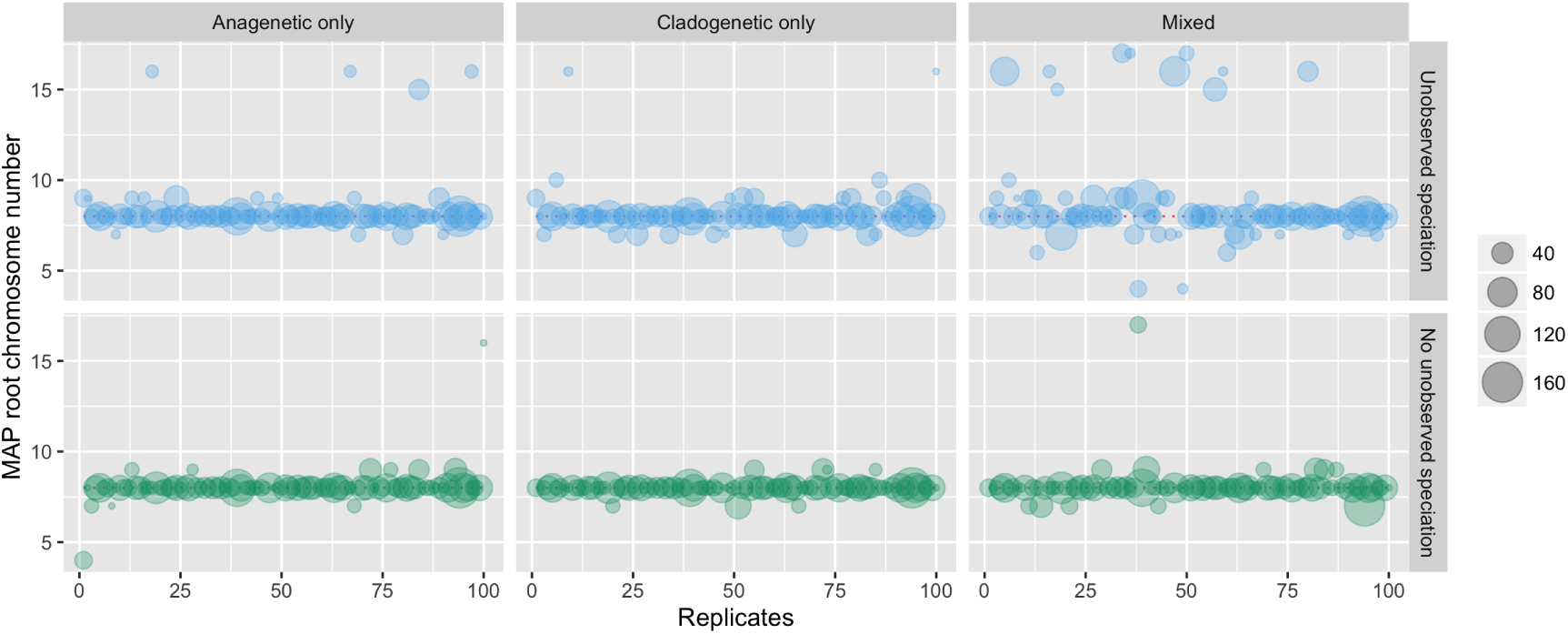
Experiment 1 results: the effect of unobserved speciation events on the maximum a posteriori (MAP) estimates of root chromosome number. Model averaged MAP estimates of the root chromosome number for 100 replicates of each simulation type on datasets that included unobserved speciation and datasets that did not include unobserved speciation. Each circle represents a simulation replicate, where the size of the circle is proportional to the number of lineages that survived to the present (the number of extant tips in the tree). The true root chromosome number used to simulate the data was 8 and is marked with a pink dotted line.

#### Experiment 2 Results

When comparing estimates from ChromoSSE that account for unobserved speciation to estimates from the non-SSE model that does not account for unobserved speciation, we found that the accuracy in estimating model parameter values was mostly similar, though for some cladogenetic parameters there was higher accuracy with the model that did account for unobserved speciation (ChromoSSE; Figure 4). For both models estimates of anagenetic parameters were more accurate than estimates of cladogenetic parameters when the true generating model included cladogenetic changes.

We found that ChromoSSE had more uncertainty in root chromosome number estimates (lower mean posterior probabilities) compared to the non-SSE model that did not account for unobserved speciation. Similarly, the root chromosome number was estimated with slightly lower accuracy (Table 4).

**Figure 4:**
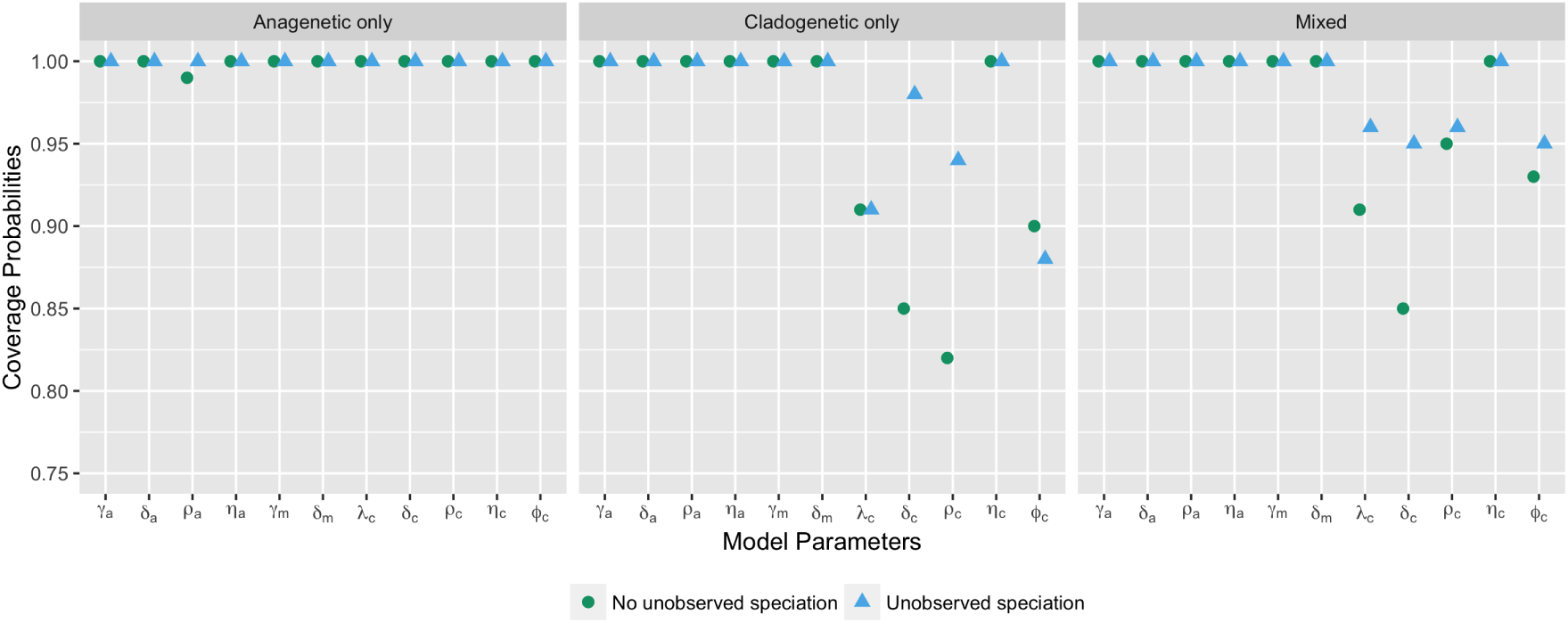
Experiment 2 results: the effect of using a model that accounts for unobserved speciation on coverage probabilities of chromosome model parameters. Each point represents the proportion of simulation replicates for which the 95% HPD interval contains the true value of the model parameter. Coverage probabilities of 1.00 mean perfect coverage. The circles represent coverage probabilities for estimates made using the non-SSE model that does not account for unobserved speciation, and the triangles represent coverage probabilities for estimates made using ChromoSSE that does account for unobserved speciation.

**Table 4:** Experiments 2 and 3 results: the effects of using a model that accounts for unobserved speciation and of jointly estimating diversification rates on ancestral chromosome number estimates. This table compares estimates of chromosome evolution using a non-SSE model that does not account for unobserved speciation events with ChromoSSE that does account for unobserved speciation events (Experiment 2), and compares estimates of chromosome evolution when jointly estimated with speciation and extinction rates versus when the true speciation and extinction rates are given (Experiment 3). Regardless of the true mode of chromosome evolution, the use of a model that accounts for unobserved speciation increases uncertainty in root state estimates. The columns from left to right are: 1) an indication of which experiment the results pertain to, 2) an indication of whether or not the estimates were made with ChromoSSE (that accounts for unobserved speciation), 3) whether diversification rates were jointly estimated with chromosome evolution, 4) the percent of simulation replicates in which the true chromosome number at the root used to simulate the data was found to be the MAP estimate, 5) the mean posterior probability of the MAP estimate of the true root chromosome number.

| Experiment # | Estimates Made w/ Model That Accounted for Unobserved Speciation? | Speciation and Extinction Rates Jointly Estimated? | Mode of Evolution Used to Simulate Data | True Root State Estimated (%) | Mean Posterior of True Root State |
| --- | --- | --- | --- | --- | --- |
| 2 | No | No | Cladogenetic | 78 | 0.87 |
| 2 | No | No | Anagenetic | 83 | 0.91 |
| 2 | No | No | Mixed | 62 | 0.80 |
| 2 & 3 | Yes | Yes | Cladogenetic | 78 | 0.81 |
| 2 & 3 | Yes | Yes | Anagenetic | 80 | 0.86 |
| 2 & 3 | Yes | Yes | Mixed | 61 | 0.72 |
| 3 | Yes | No | Cladogenetic | 78 | 0.84 |
| 3 | Yes | No | Anagenetic | 83 | 0.90 |
| 3 | Yes | No | Mixed | 62 | 0.76 |

#### Experiment 3 Results

We found that jointly estimating speciation and extinction rates with chromosome number evolution using ChromoSSE slightly decreased the accuracy of root chromosome number estimates, and further it increased the uncertainty of the inferred root chromosome number (as reflected in lower mean posterior probabilities; Table 4). Fixing the speciation and extinction rates to their true value removed much of the increased uncertainty associated with using a model that accounts for unobserved speciation (Table 4).

#### Experiment 4 Results

Under simulation scenarios that had cladogenetic changes but no anagenetic changes, we found that ChromoSSE overestimated anagenetic parameters and underestimated cladogenetic parameters (Figure 5A), which explains the lower coverage probabilities of cladogenetic parameters reported above for experiment 2 (Figure 4). When anagenetic parameters were fixed to 0.0 cladogenetic parameters were no longer underestimated (Figure 5 A), and the coverage probabilities of cladogenetic parameters increased slightly (Figure 5 B).

**Figure 5:**
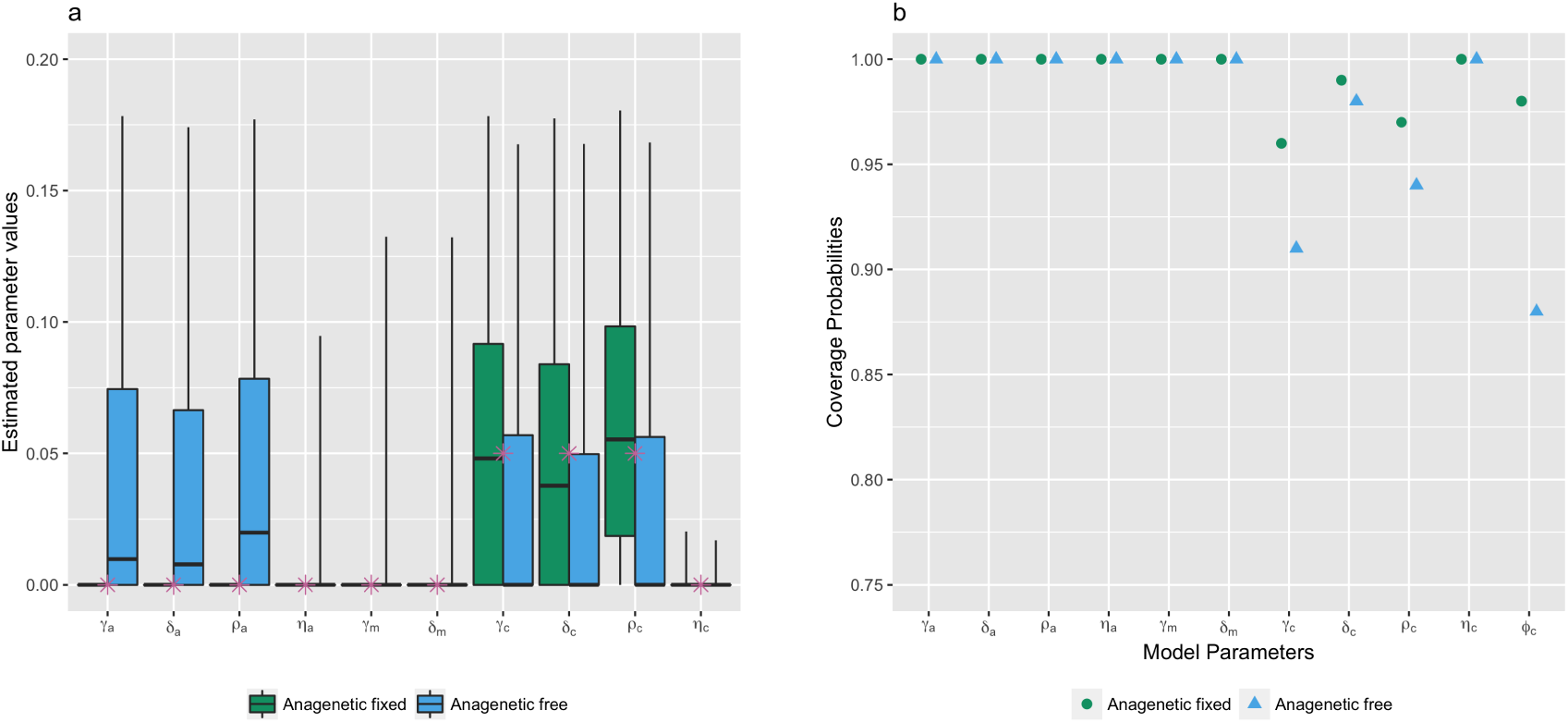
Experiment 4 results: testing identifiability of cladogenetic parameters under ChromoSSE. a) Chromosome parameter value estimates from 100 simulation replicates under a simulation scenario with no anagenetic changes (cladogenetic only). The stars represent true values. The box plots compare parameter estimates made when anagenetic parameters were fixed to 0 to estimates made when all parameters were free. When all parameters were free the anagenetic parameters were overestimated and cladogenetic parameters were underestimated. When the anagenetic parameters were fixed to 0 the estimates for the cladogenetic parameters were more accurate. b) Coverage probabilities of chromosome evolution parameters under the cladogenetic only model of chromosome evolution. The accuracy of cladogenetic parameter estimates increased when anagenetic parameters were fixed to 0.

#### Experiment 5 Results

We found that incomplete sampling of extant lineages had a minor effect on the accuracy of ancestral chromosome number estimates (Figure 6). Accuracy only slightly decreased as the percentage of extant lineages sampled declined from 100% to 50%, and decreased more rapidly when the percentage went to 10%. As measured by the proportion of simulation replicates that inferred the MAP root chromosome number to be the true root chromosome number, the accuracy of ancestral states estimated under ChromoSSE declined from 0.80 accuracy at 100% taxon sampling to 0.69 at 10% taxon sampling. Essentially no difference in accuracy was detected between the non-SSE model that does not take unobserved speciation into account and ChromoSSE. Furthermore, little difference in accuracy was detected using ChromoSSE with the taxon sampling probability *ρ_s_* set to 1.0 compared to ChromoSSE with *ρ_s_* set to the true value (0.1, 0.5, or 1.0; Figure 6).

**Figure 6:**
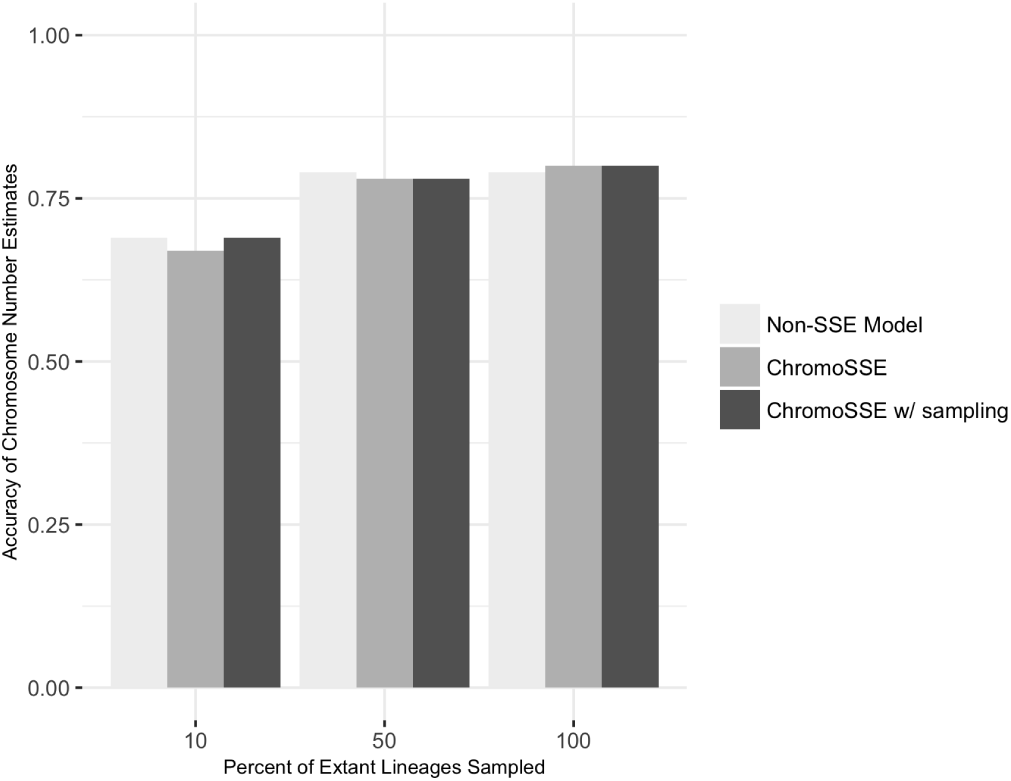
Experiment 5 results: the effect of incomplete sampling. The accuracy of ancestral chromosome number estimates slightly declined as the percentage of sampled extant lineages decreased from 100% to 50%, and decreased more quickly once the percentage of extant lineages decreased to 10%. There was little difference between the non-SSE model (light grey) that does not take into account unobserved speciation and ChromoSSE (medium and dark grey) which does take into account unobserved speciation. Furthermore, little difference in accuracy was detected using ChromoSSE with the taxon sampling probability *ρ_s_* set to 1.0 (medium grey) and with *ρ_s_* set to the true value (0.1, 0.5, or 1.0; dark grey). The accuracy of chromosome number estimates was measured by the proportion of simulation replicates for which the estimated MAP root chromosome number corresponded with the true chromosome number used to simulate the data.

### Empirical Data

Model averaged MAP estimates of ancestral chromosome numbers for each of the five empirical datasets are show in Figures 7, 8, 9, 10, and 11. The mean model-averaged chromosome number evolution parameter value estimates for the empirical datasets are reported in Table 5. Posterior probabilities for the MAP model of chromosome number evolution were low for all datasets, varying between 0.04 for *Carex* section *Spirostachyae* and 0.21 for *Helianthus* (Table 6). Bayes factors supported unique, clade-specific combinations of anagenetic and cladogenetic parameters for all five datasets (Table 6). None of the clades had support for purely anagenetic or purely cladogenetic models of chromosome evolution.

**Figure 7:**
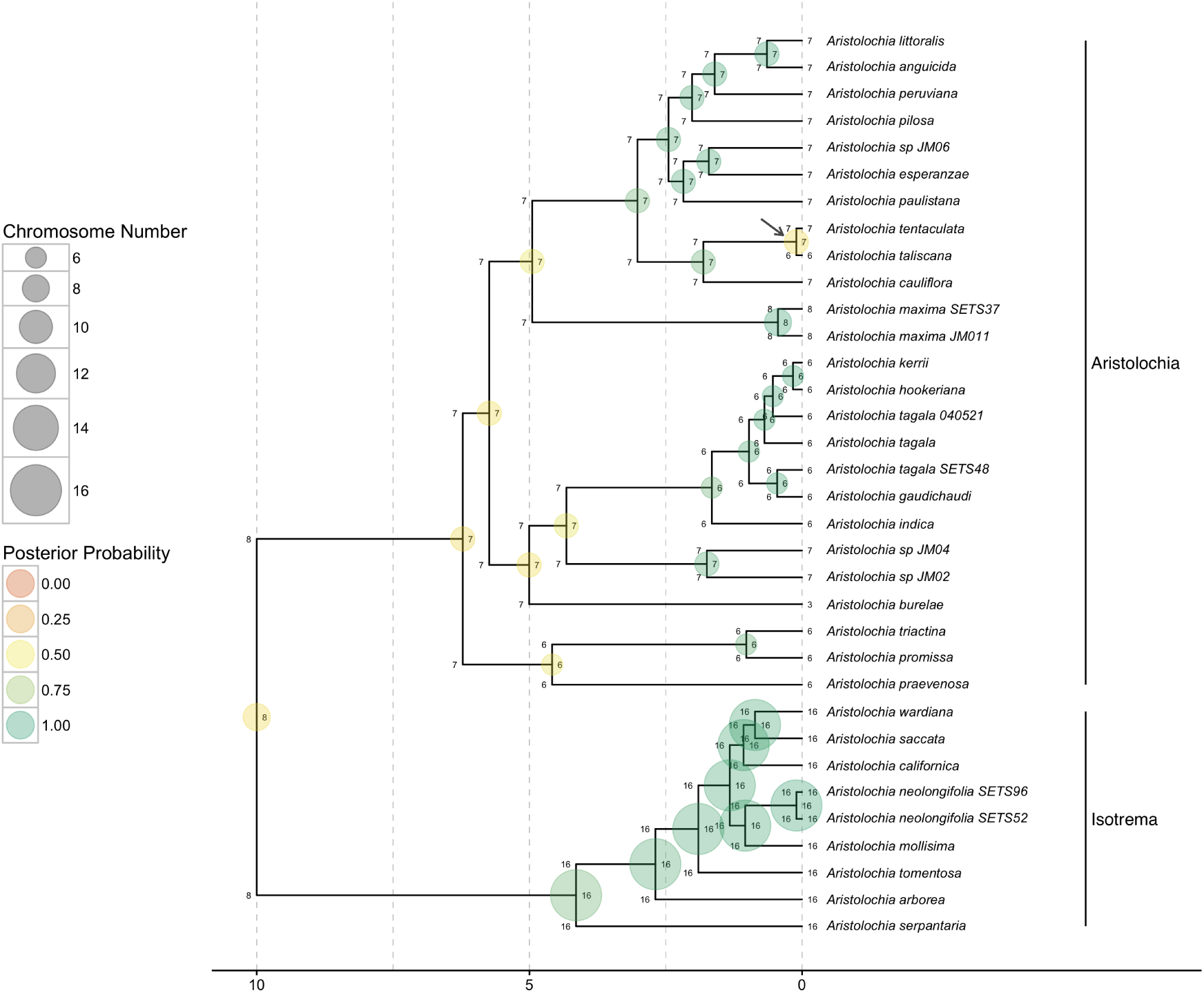
Ancestral chromosome number estimates of *Aristolochia*. The model averaged MAP estimate of ancestral chromosome numbers are shown at each branch node. The states of each daughter lineage immediately after cladogenesis are shown at the “shoulders” of each node. The size of each circle is proportional to the chromosome number and the color represents the posterior probability. The MAP root chromosome number is 8 with a posterior probability of 0.45. The grey arrow highlights the possible dysploid speciation event leading to the west-central Mexican species *Aristolochia tentaculata* and *A. taliscana*. Clades corresponding to subgenera are indicated at right.

**Figure 8:**
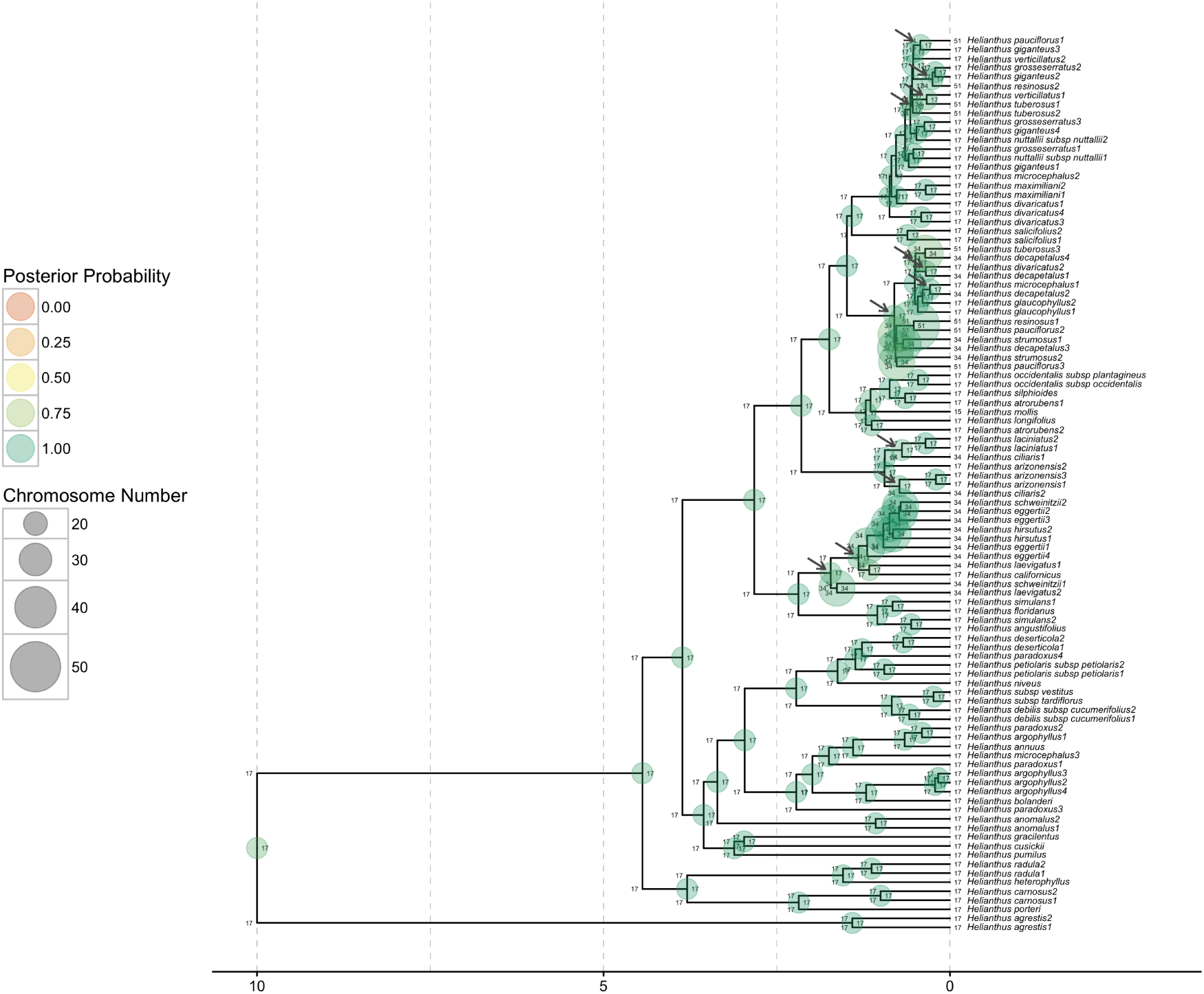
Ancestral chromosome number estimates of *Helianthus*. The model averaged MAP estimate of ancestral chromosome numbers are shown at each branch node. The states of each daughter lineage immediately after cladogenesis are shown at the “shoulders” of each node. The size of each circle is proportional to the chromosome number and the color represents the posterior probability. The MAP root chromosome number is 17 with a posterior probability of 0.91. The grey arrows show the locations of 12 inferred polyploid speciation events.

**Figure 9:**
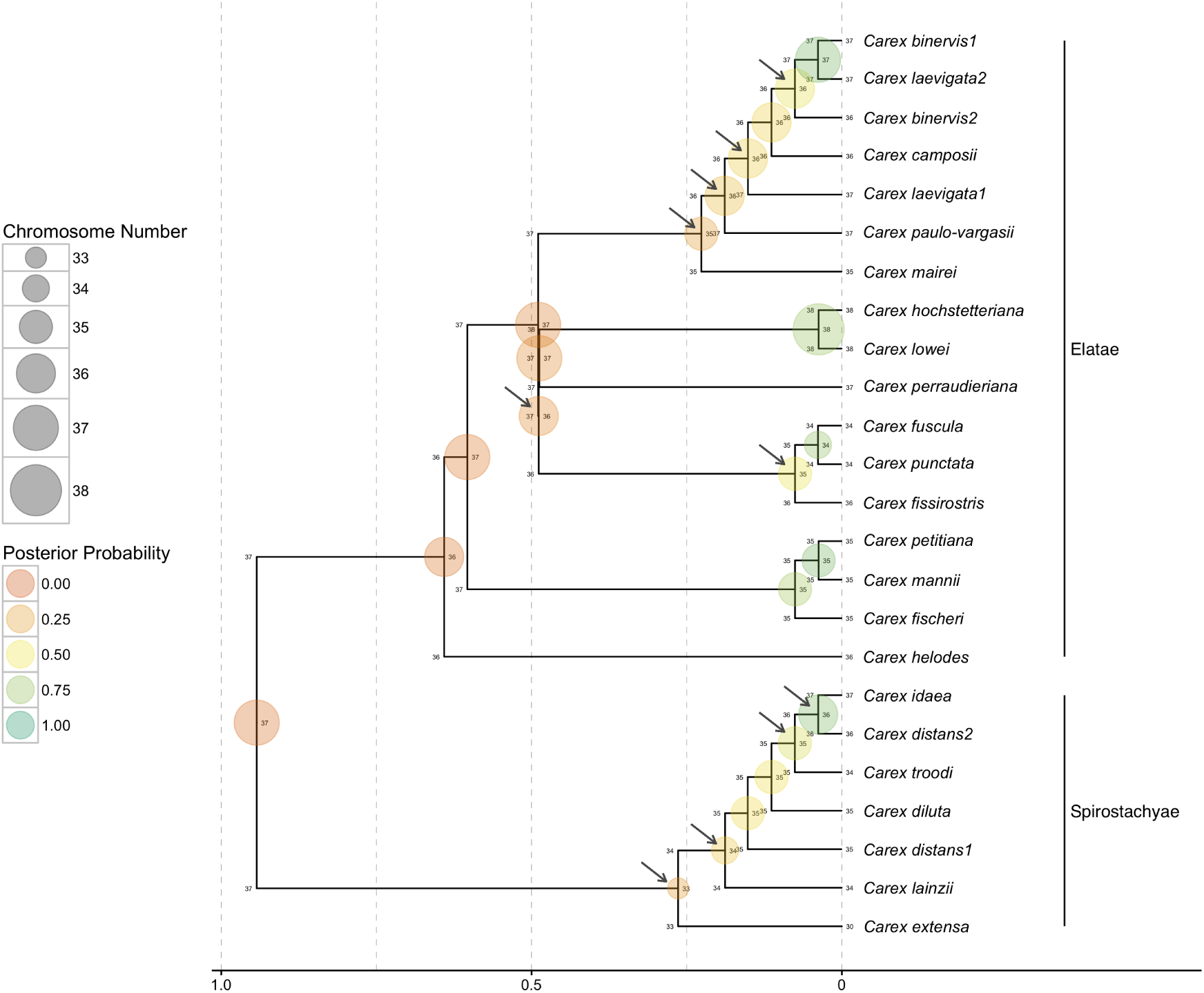
Ancestral chromosome number estimates of *Carex* section *Spirostachyae*. The model averaged MAP estimate of ancestral chromosome numbers are shown at each branch node. The states of each daughter lineage immediately after cladogenesis are shown at the “shoulders” of each node. The size of each circle is proportional to the chromosome number and the color represents the posterior probability. The MAP root chromosome number is 37 with a posterior probability of 0.08. Grey arrows indicate the location of possible dysploid speciation events. 36.9% of all speciation events include a cladogenetic gain or loss of a single chromosome. Clades corresponding to subsections are indicated at right.

**Figure 10:**
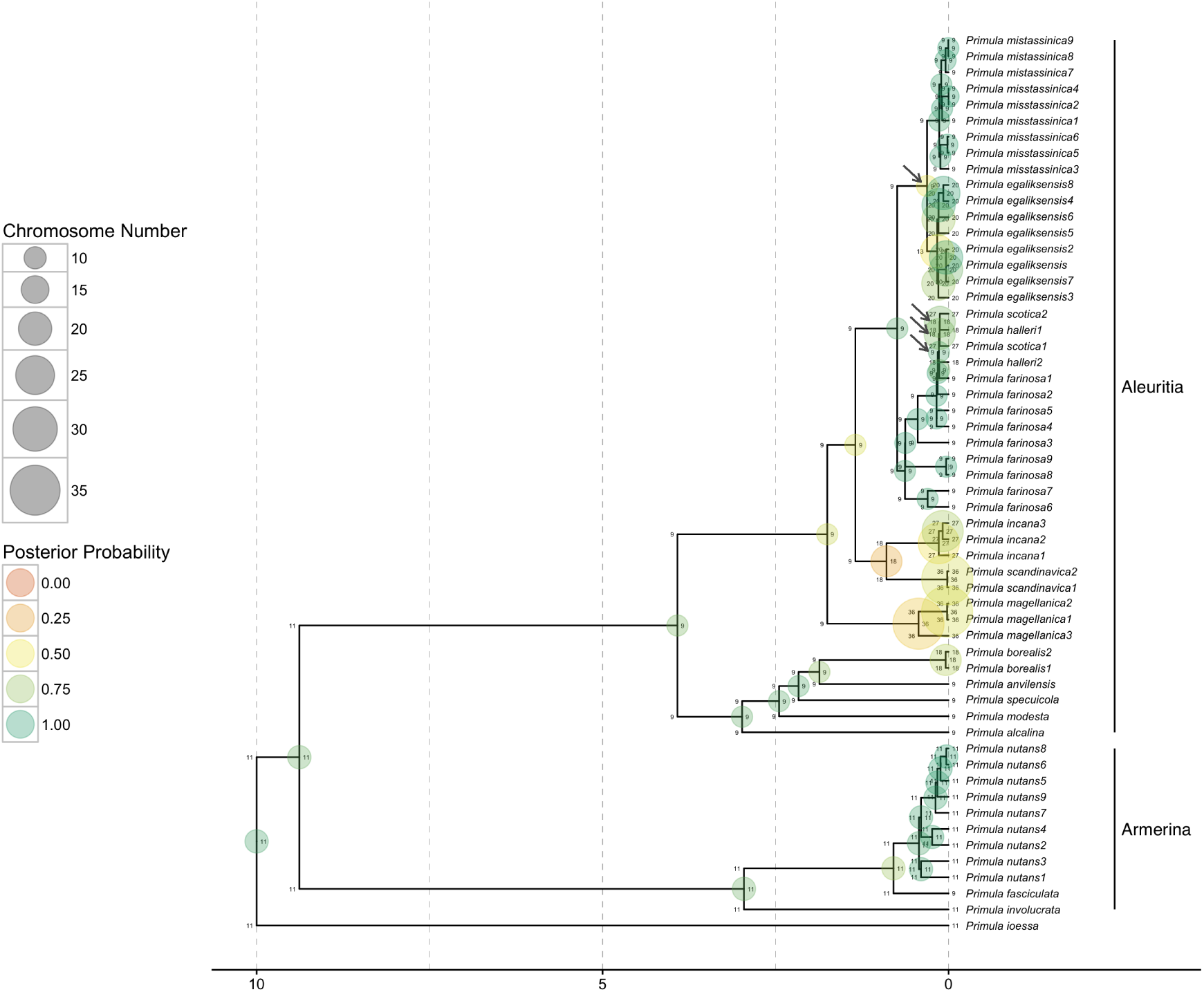
Ancestral chromosome number estimates of *Primula* section *Aleuritia*. The model averaged MAP estimate of ancestral chromosome numbers are shown at each branch node. The states of each daughter lineage immediately after cladogenesis are shown at the “shoulders” of each node. The size of each circle is proportional to the chromosome number and the color represents the posterior probability. The MAP root chromosome number of section *Aleuritia* is 9 with a posterior probability of 0.82. The arrows show the inferred history of possible polyploid and demi-polyploid speciation events in the clade containing the tetraploids *Primula egaliksensis* and *P. halleri* and the hexaploid *P. scotica*. Clades corresponding to sections are indicated at right.

**Figure 11:**
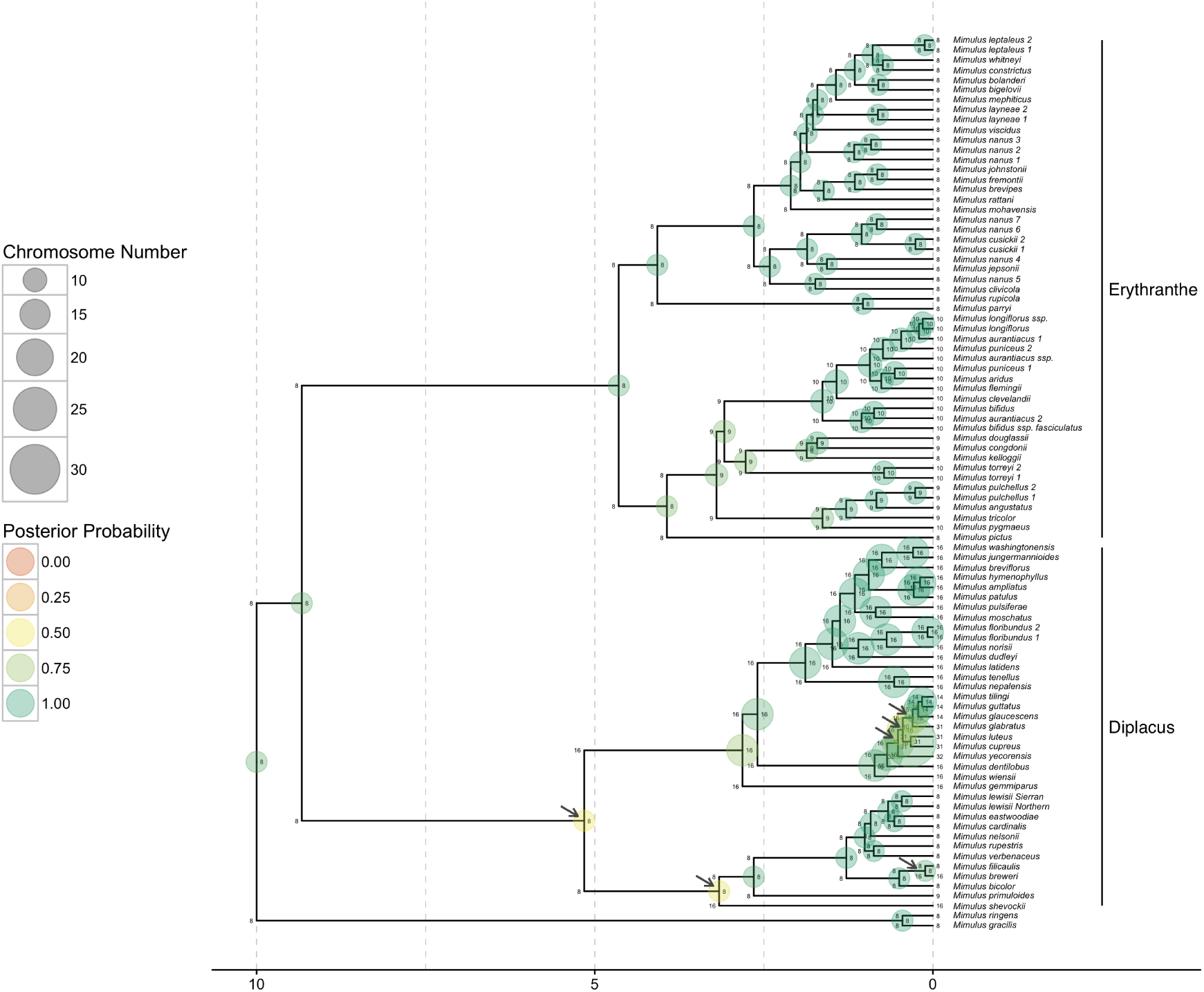
Ancestral chromosome number estimates of *Mimulus* sensu lato. The model averaged MAP estimate of ancestral chromosome numbers are shown at each branch node. The states of each daughter lineage immediately after cladogenesis are shown at the “shoulders” of each node. The size of each circle is proportional to the chromosome number and the color represents the posterior probability. The MAP root chromosome number is 8 with a posterior probability of 0.90. The arrows highlight the inferred history of repeated polyploid speciation events in the Diplacus clade, which contains the tetraploids *Mimulus cupreus, M. glabratus, M. luteus*, and *M. yecorensis*. Clades corresponding to segregate genera are indicated at right.

**Table 5:** Mean model-averaged parameter value estimates for empirical datasets. Rates for all parameters are given in units of chromosome changes per branch length unit except for μ which is given in extinction events per time units.

| Clade | $\gamma_a$ | $\delta_a$ | $\rho_a$ | $\eta_a$ | $\gamma_m$ | $\delta_m$ | $\phi_c$ | $\gamma_c$ | $\delta_c$ | $\rho_c$ | $\eta_c$ | $\mu$ |
| --- | --- | --- | --- | --- | --- | --- | --- | --- | --- | --- | --- | --- |
| *Aristolochia* | 0.02 | 0.05 | 0.01 | 0.0 | -0.01 | -0.01 | 0.43 | 0.0 | 0.04 | 0.0 | 0.0 | 0.19 |
| *Carex* section<br>*Spirostachyae* | 0.19 | 0.79 | 0.16 | 0.13 | 0.0 | 0.04 | 2.49 | 2.15 | 0.15 | 0.95 | 0.5 | 2.26 |
| *Helianthus* | 0.0 | 0.02 | 0.0 | 0.03 | -0.0 | -0.0 | 0.68 | 0.0 | 0.0 | 0.13 | 0.0 | 0.09 |
| *Mimulus* s.l. | 0.03 | 0.02 | 0.01 | 0.0 | 0.02 | 0.02 | 0.65 | 0.0 | 0.0 | 0.05 | 0.0 | 0.16 |
| *Primula*<br>section<br>*Aleuritia* | 0.01 | 0.05 | 0.01 | 0.01 | -0.0 | -0.0 | 2.39 | 0.01 | 0.03 | 0.15 | 0.09 | 2.47 |

**Table 6:**
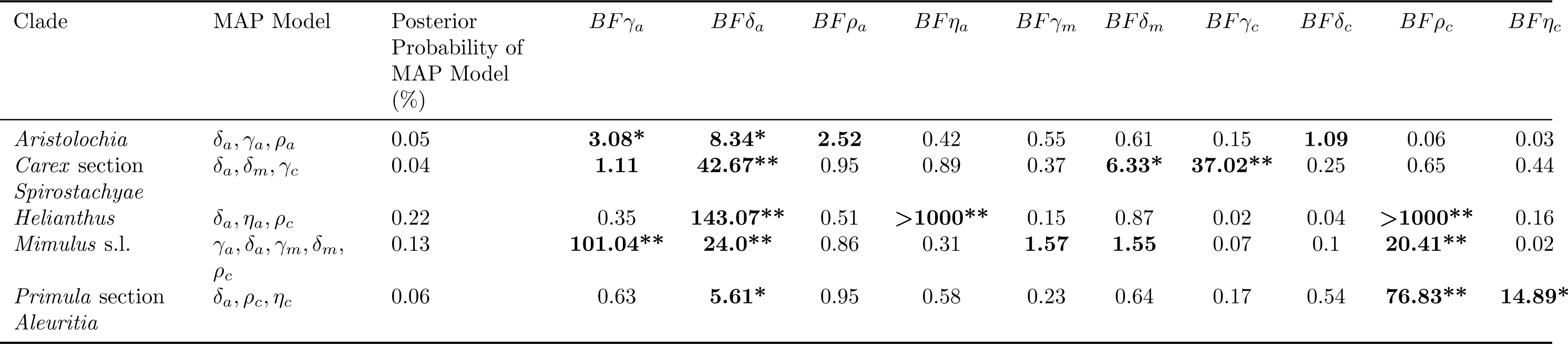
Best supported chromosome evolution models for empirical datasets. The MAP model of chromosome evolution and its corresponding posterior probability are shown with Bayes factors (*BF*) for models that include each parameter. Parameters with *BF* > 1 are in bold and indicate support for models that include that parameter. Parameters with “positive” and “strong” support according to Kass and Raftery (1995) are marked with * and **, respectively.

The ancestral state reconstructions for *Aristolochia* were highly similar to those found by Mayrose et al. (2010). We found a moderately supported root chromosome number of 8 (posterior probability 0.45), and a polyploidization event on the branch leading to the *Isotrema* clade which has a base chromosome number of 16 with high posterior probability (0.88; Figure 7). On the branch leading to the main *Aristolochia* clade we found a dysploid loss of a single chromosome. Overall, we estimated moderate rates of anagenetic dysploid and polyploid changes, and the rates of cladogenetic change were 0 except for a moderate rate of cladogenetic dysploid loss (Tables 5). There was only one cladogenetic change inferred in the MAP ancestral state reconstruction, which was a recent possible dysploid speciation event that split the sympatric west-central Mexican species *Aristolochia tentaculata* and *A. taliscana*.

In *Helianthus*, on the other hand, we found high rates of cladogenetic polyploidization, and low rates of anagenetic change (Tables 5). 12 separate possible polyploid speciation events were identified over the phylogeny (Figure 8), and cladogenetic polyploidization made up 16% of all observed and unobserved speciation events. Bayes factors gave very strong support for models that included cladogenetic polyploidization as well as anagenetic demi-polyploidization (Table 6), the latter explaining the frequent anagenetic transitions from 34 to 51 chromosomes found in the MAP ancestral state reconstruction. The well supported root chromosome number of 17 (posterior probability 0.91) corresponded with the findings of Mayrose et al. (2010).

As opposed to the *Helianthus* results, the *Carex* section *Spirostachyae* estimates had very low rates of polyploidization and instead had high rates of cladogenetic dysploid change (Tables 5). An estimated 36.9% of all observed and unobserved speciation events included a cladogenetic gain or loss of a single chromosome. Overall, the rates of anagenetic changes were estimated to be much lower than the rates of cladogenetic changes. Bayes factors did not support either anagenetic or cladogenetic polyploidization (Table 6). The MAP root chromosome number of 37, despite being very weakly supported (0.08), corresponds with the findings of Escudero et al. (2014), where it was also poorly supported (Figure 9).

In *Primula*, we found a base chromosome number for section *Aleuritia* of 9 with high posterior probability (0.82; Figure 10), which agrees with estimates from Glick and Mayrose (2014). We estimated moderate rates of anagenetic and cladogenetic changes, including both cladogenetic polyploidization and demi-polyploidization (Table 5). The MAP ancestral state estimates include an inferred history of possible polyploid and demi-polyploid speciation events in the clade containing the tetraploid *Primula halleri* and the hexaploid *P. scotica. Primula* is the only dataset out of the five analysed here for which Bayes factors supported the inclusion of cladogenetic demi-polyploidization (Table 6).

The well supported root chromosome number of 8 (posterior probability 0.90) found for *Mimulus* s.l. corresponds with the inferences reported in Beardsley et al. (2004). We estimated moderate rates of anagenetic dysploid gains and losses, as well as a moderate rate of cladogenetic polyploidization (Table 5). Bayes factors also supported models that included anagenetic dysploid gain and loss, as well as cladogenetic polyploidization (Table 6). The MAP ancestral state reconstruction revealed that most of the possible polyploid speciation events took place in the *Diplacus* clade, particularly in the clade containing the tetraploids *Mimulus cupreus, M. glabratus, M. luteus*, and *M. yecorensis* (Figure 11). Additionally, an ancient cladogenetic polyploidization event is inferred for the split between the two main *Diplacus* clades at about 5 million time units ago.

## Discussion

The results from the empirical analyses show that the ChromoSSE models detect strikingly different modes of chromosome evolution with clade-specific combinations of anagenetic and cladogenetic processes. Anagenetic dysploid gains and losses were supported in nearly all clades; however, cladogenetic dysploid changes were supported only in *Carex*. The occurrence of anagenetic dysploid changes in all clades suggest that small chromosome number changes due to gains and losses may frequently have a minimal effect on the formation of reproductive isolation, though our results suggest that *Carex* may be a notable exception. Anagenetic polyploidization was only supported in *Aristolochia*, while cladogenetic polyploidization was supported in *Helianthus, Mimulus* s.l., and *Primula*. These findings confirm the evidence presented by Zhan et al. (2016) that polyploidization events could play a significant role during plant speciation.

Our models shed new light on the importance of whole genome duplications as a key driver in evolutionary diversification processes. *Helianthus* has long been understood to have a complex history of polyploid speciation (Timme et al. 2007), but our results here are the first to statistically show the prevalance of cladogenetic polyploidization in *Helianthus* (occuring at 16% of all speciation events) and how few of the chromosome changes are estimated to be anagenetic. Polyploid speciation has also been suspected to be common in *Mimulus* s.l. (Vickery 1995), and indeed we estimated that 7% of speciation events were cladogenetic polyploidization events. We also estimated that the rates of cladogenetic dysploidization in *Mimulus* s.l. were 0, which is in contrast to the parsimony based inferences presented in Beardsley et al. (2004), which estimated 11.5% of all speciation events included polyploidization and 13.3% included dysploidization. Their estimates, however, did not distinguish cladogenetic from anagenetic processes, and so they likely underestimated anagenetic changes. Our ancestral state reconstructions of chromosome number evolution for *Helianthus, Mimulus* s.l., and *Primula* show that polyploidization events generally occurred in the relatively recent past; few ancient polyploidization events were reconstructed (one exception being the ancient cladogenetic polyploidization event in *Mimulus* clade *Diplacus*). This pattern appears to be consistent with recent studies that show polyploid lineages may undergo decreased net diversification (Mayrose et al. 2011; Scarpino et al. 2014), leading some to suggest that polyploidization may be an evolutionary dead-end (Arrigo and Barker 2012). While in the analyses presented here we fixed rates of speciation and extinction through time and across lineages, an obvious extension of our models would be to allow these rates to vary across the tree and statistically test for rate changes in polyploid lineages.

Our findings also suggest dysploid changes may play a significant role in the speciation process of some lineages. The genus *Carex* is distinguished by holocentric chromosomes that undergo common fusion and fission events but rarely polyploidization (Hipp 2007). This concurs with our findings from *Carex* section *Spirostachyae*, where we saw no support for models including either anagenetic or cladogenetic polyploidization. Instead we found high rates of cladogenetic dysploid change, which is congruent with earlier results that show that *Carex* diversification is driven by processes of fission and fusion occurring with cladogenetic shifts in chromosome number (Hipp 2007; Hipp et al. 2007). Hipp (2007) proposed a speciation scenario for *Carex* in which the gradual accumulation of chromosome fusions, fissions, and rearrangements in recently diverged populations increasingly reduce the fertility of hybrids between populations, resulting in high species richness. More recently, Escudero et al. (2016) found that chromosome number differences in *Carex scoparia* led to reduced germination rates, suggesting hybrid dysfunction could spur chromosome speciation in *Carex*. Holocentricity has arisen at least 13 times independently in plants and animals (Melters et al. 2012), thus future work could examine chromosome number evolution in other holocentric clades and test for similar patterns of cladogenetic fission and fusion events.

The models presented here could also be used to further study the role of divergence in genomic architecture during sympatric speciation. Chromosome structural differences have been proposed to perform a central role in sympatric speciation, both in plants (Gottlieb 1973) and animals (Feder et al. 2005; Michel et al. 2010). In *Aristolochia* we found most changes in chromosome number were estimated to be anagenetic, with the only cladogenetic change occuring among a pair of recently diverged sympatric species. By coupling our chromosome evolution models with models of geographic range evolution it would be possible to statistically test whether the frequency of cladogenetic chromosome changes increase in sympatric speciation events compared to allopatric speciation events, thereby testing for interaction between these two different processes of reproductive isolation and evolutionary divergence.

The simulation results from Experiment 1 demonstrate that extinction reduces the accuracy of inferences made by models of chromosome evolution that do not take into account unobserved speciation events. Furthermore, the simulations performed in Experiments 2 and 3 show that the substantial uncertainty introduced in our analyses by jointly estimating diversification rates and chromosome evolution resulted in lower posterior probabilities for ancestral state reconstructions. We feel that this is a strength of our method; the lower posterior probabilities incorporate true uncertainty due to extinction and so represent more conservative estimates. Additionally, the simulation results from Experiment 4 reveal that rates of anagenetic evolution were overestimated and rates of cladogenetic change were underestimated when the generating process consisted only of cladogenetic events. This suggests the possibility that our models of chromosome number evolution are only partially identifiable, and that the results of our empirical analyses may have a similar bias towards overestimating anagenetic evolution and underestimating cladogenetic evolution. This bias may be an issue for all ClaSSE type models, but the practical consequences here are conservative estimates of cladogenetic chromosome evolution.

An important caveat for all phylogenetic methods is that estimates of model parameters and ancestral states can be highly sensitive to taxon sampling (Heath et al. 2008). All of the empirical datasets examined here included non-monophyletic taxa that were treated as separate lineages. We made the unrealistic assumptions that 1) each of the non-monophyletic lineages sharing a taxon name have the same cytotype, and 2) the taxon sampling probability (*ρ_s_*) for the birth-death process was 1.0. The former assumption could drastically affect ancestral state estimates, but its effect can only be confirmed by obtaining chromosome counts for each lineage regardless of taxon name. While the results from simulation Experiment 5 showed that fixing *ρ_s_* to 1.0 did not decrease the accuracy of inferred ancestral states, we still performed extra analyses of the empirical datasets with different values of *ρ_s_* (results not shown). The results indicated that total speciation and extinction rates are sensitive to *ρ_s_*, but the relative speciation rates (e.g. between *ϕ_c_* and *γ_c_*) remained similar. The ancestral state estimates of cladogenetic and anagenetic chromosome changes were robust to different values of *ρ_s_*. This could vary among datasets and care should be taken when considering which lineages to sample.

Bayesian model averaging is particularly appropriate for models of chromosome number evolution since conditioning on a single model ignores the considerable degree of model uncertainty found in both the simulations and the empirical analyses. In the simulations the true model of chromosome evolution was rarely inferred to be the MAP model (< 39% of replicates), and in the instances it was correctly identified the posterior probability of the MAP model was < 0.13. The posterior probabilities of the MAP models for the empirical datasets were similarly low, varying between 0.04 and 0.22. Conditioning on a single poorly fitting model of chromosome evolution, even when it is the best model available, results in an underestimate of the uncertainty of ancestral chromosome numbers. Furthermore, Bayesian model averaging enabled us to detect different modes of chromosome number evolution without the limitation of traditional model testing procedures in which multiple analyses are performed that each condition on a different single model. This is a particularly useful approach when the space of all possible models is large.

Our RevBayes implementation facilitates model modularity and easy experimentation. Experimenting with different priors or MCMC moves is achieved by simply editing the Rev scripts that describe the model. Though in our analyses here we ignored phylogenetic uncertainty by assuming a fixed known tree, we could easily incorporate this uncertainty by modifying a couple lines of the Rev script to integrate over a previously estimated posterior distribution of trees. We could also use molecular sequence data simultaneously with the chromosome models to jointly infer phylogeny and chromosome evolution, allowing the chromosome data to help inform tree topology and divergence times. In this paper we chose not to perform joint inference so that we could isolate the behavior of the chromosome evolution models; however, this is a promising direction for future research.

There are a number of challenging directions for future work on phylogenetic chromosome evolution models. Models that incorporate multiple aspects of chromosome morphology such as translocations, inversions, and other gene synteny data as well as the presence of ring and/or B chromosomes have yet to be developed. None of our models currently account for allopolyploidization; indeed few phylogenetic comparative methods can handle reticulate evolutionary scenarios that result from allopolyploidization and other forms of hybridization (Marcussen et al. 2015). A more tractable problem is mapping chromosome number changes along the branches of the phylogeny, as opposed to simply making estimates at the nodes as we have done here. Since the approach described here models both anagenetic and cladogenetic chromosome evolution processes while accounting for unobserved spéciation events, the rejection sampling procedure used in standard stochastic character mapping (Nielsen 2002; Huelsenbeck et al. 2003) is not sufficient. While data augmentation approaches such as those described by Bokma (2008) could be utilized, they require complex MCMC algorithms that may have difficulty mixing. Another option is to extend the method described in this paper to draw joint ancestral states by numerically integrating root-to-tip over the tree into a new procedure called joint conditional character mapping. This sort of approach would infer the joint MAP history of chromosome changes both at the nodes and along the branches of the tree, and provide an alternative to stochastic character mapping that will work for all ClaSSE type models.

### Conclusions

The analyses presented here show that the ChromoSSE models of chromosome number evolution successfully infer different clade-specific modes of chromosome evolution as well as the history of anagenetic and cladogenetic chromosome number changes for a clade, including reconstructing the timing and location of possible chromosome speciation events over the phylogeny. These models will help investigators study the mode and history of chromosome evolution within individual clades of interest as well as advance understanding of how fundamental changes in the architecture of the genome such as whole genome duplications affect macroevolutionary patterns and processes across the tree of life.

## Acknowledgements

Thank you to Bruce Baldwin, Emma Goldberg, and Michael Landis for valuable discussions. We also wish to thank two anonymous reviewers for their thoughtful feedback that improved this work.

## Supplementary Material

## Appendix 1: Validating RevBayes Ancestral State Estimates

### Ancestral State Estimates of SSE Models

The code repository http://github.com/wf8/anc_state_validation contains scripts to validate the Monte Carlo method of ancestral state estimation for state-dependent speciation and extinction (SSE) models we implemented in RevBayes (Höhna et al. 2016) against the analytical marginal ancestral state estimation implemented in the R package diversitree (FitzJohn 2012).

Although the closest model to ChromoSSE implemented in diversitree is ClaSSE (Goldberg and Igić 2012), ancestral state estimation for ClaSSE is not implemented in diversitree. Therefore here we compare the ancestral state estimates for BiSSE (Maddison et al. 2007) as implemented in diversitree to the estimates made by RevBayes. Note that as implemented in RevBayes the BiSSE, ChromoSSE, ClaSSE, MuSSE (FitzJohn 2012), and HiSSE (Beaulieu and O’Meara 2016) models use the same C++ classes and algorithms for parameter and ancestral state estimation, so validating ancestral state estimates for BiSSE should provide confidence in estimates made by RevBayes for all these SSE models.

In RevBayes we sample ancestral states for SSE models from their joint distribution conditional on the tip states and the model parameters during the MCMC. However, in this work we summarize the MCMC samples by calculating the marginal posterior probability of each node being in each state. So the RevBayes marginal ancestral state reconstructions which are estimated via MCMC are directly comparable to the analytical marginal ancestral states computed by diversitree. It would be possible to summarize the samples from the MCMC to reconstruct the maximum a posteriori joint ancestral state reconstruction, but we have not done so in this work.

### Comparison of RevBayes Estimates to Diversitree

Here we show ancestral state estimates under BiSSE for an example where the tree and tip data were simulated in diversitree with the following parameters: *λ*_0_ = 0.2, *λ*_1_ = 0.4,*μ*_0_ = 0.01, *μ*_1_ = 0.1, and *q*_01_ = *q*_10_ = 0.1. The ancestral state reconstructions from RevBayes and diversitree are shown in Figures 2 and 3, respectively.

**Figure 1:**
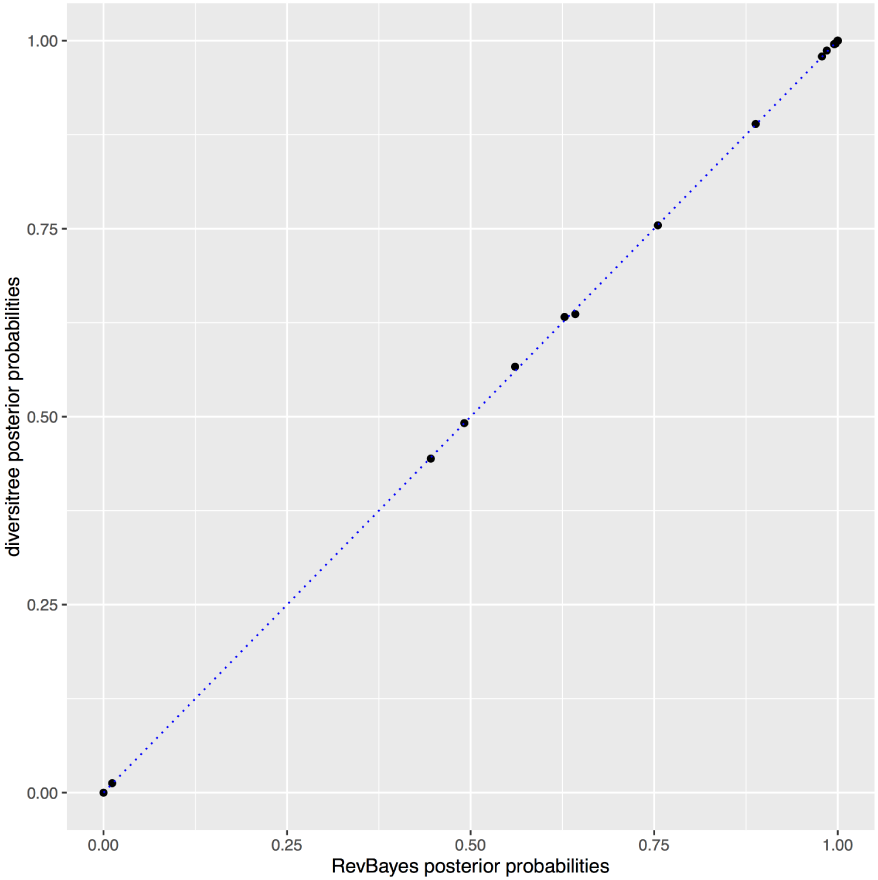
Posterior probabilities of marginal ancestral state estimates. Each point represents the marginal posterior probability of a node being in state 1 as estimated by RevBayes plotted against the estimates made by diversitree. The marginal ancestral states were estimated under BiSSE from a tree and tip data simulated with the following parameters: *λ*_0_ = 0.2, *λ*_1_ = 0.4, *μ*_0_ = 0.01, *μ*_1_ = 0.1, and *q*_01_ = *q*_10_ = 0.1. The full ancestral state reconstructions from RevBayes and diversitree are shown in Figures 2 and 3, respectively.

**Figure 2:**
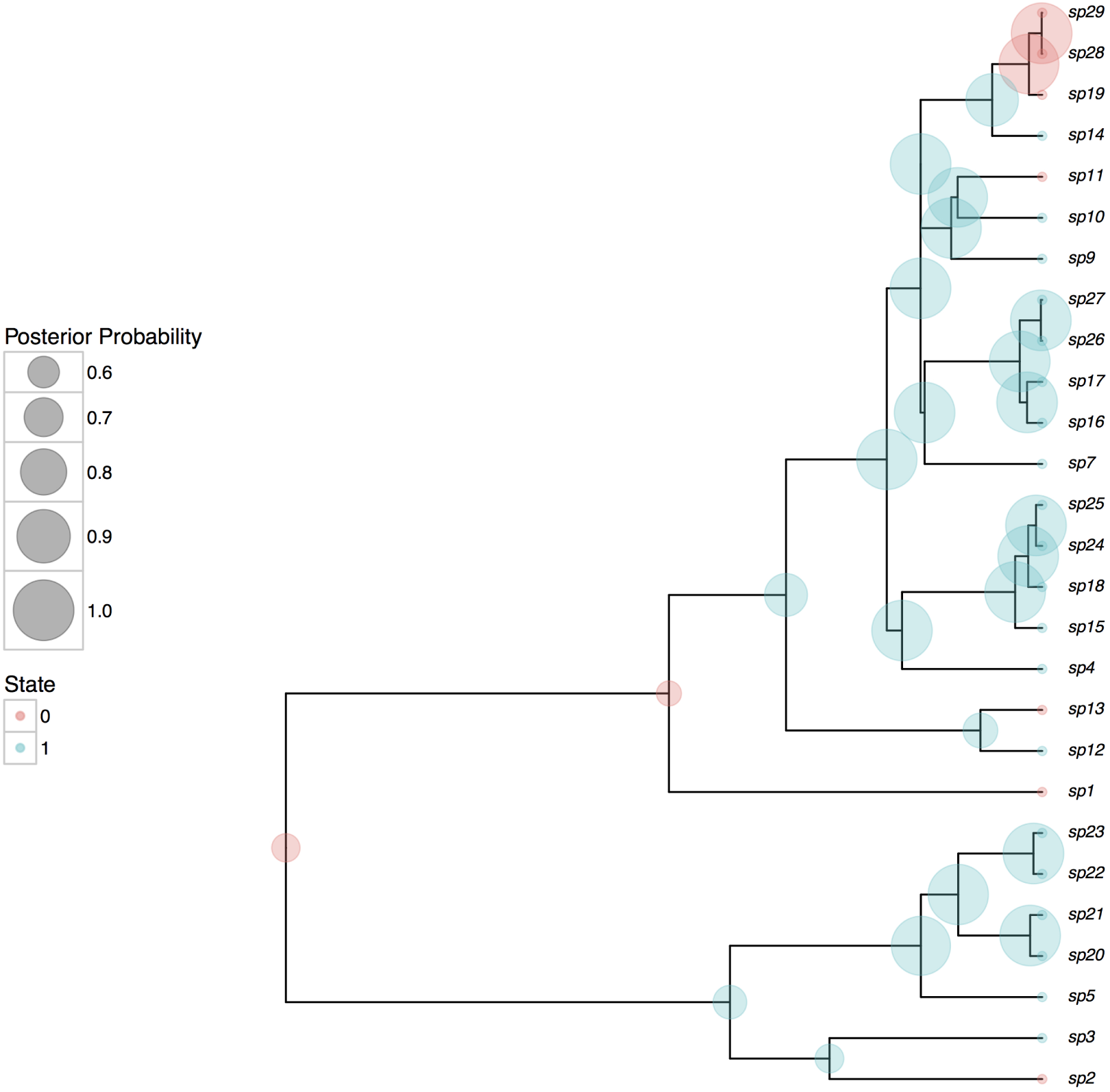
Ancestral state estimates from RevBayes. Marginal ancestral states estimated under BiSSE from a tree and tip data simulated with the following parameters: *λ*_0_ = 0.2, *λ*_1_ = 0.4, *μ*_0_ = 0.01, *μ*_1_ = 0.1, and *q*_01_ = *q*_10_ = 0.1.

**Figure 3:**
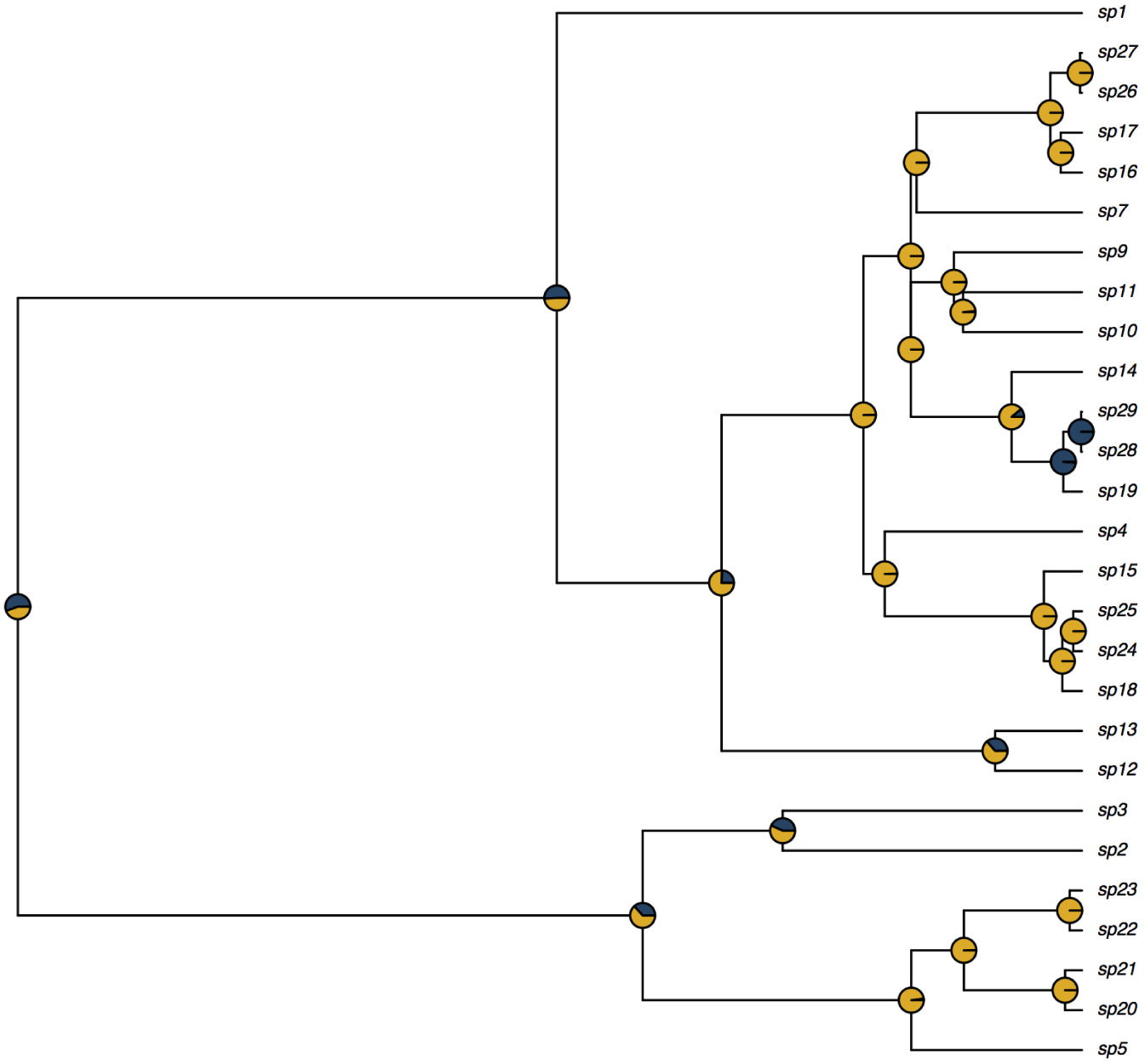
Ancestral state estimates from diversitree. Marginal ancestral states estimated under BiSSE from a tree and tip data simulated with the following parameters: *λ*_0_ = 0.2, *λ*_1_ = 0.4, *μ*_0_ = 0.01, *μ*_1_ = 0.1, and *q*_01_ = *q*_10_ = 0.1.

The log-likelihood as computed by diversitree was -109.46, whereas with RevBayes it was -109.71. Small differences in the log-likelihoods are expected due to differences in the way diversitree and RevBayes calculate probabilities at the root, and also due to numerical approximations. However both reconstructions should return the same probabilities for ancestral states at the root, and indeed diversitree calculated the root probability of being in state 0 as 0.555 and RevBayes calculated it as 0.554. The estimated posterior probabilities are very close for all nodes. This is shown in a plot comparing the marginal posterior probabilities for all nodes being in state 1 as estimated by RevBayes against the diversitree estimates (Figure 1).

## Appendix 2: Metropolis-Hastings Moves

The Metropolis-Hastings moves used in all ChromoSSE analyses are outlined in Table 1. All MCMC proposals are standard except the ElementSwapSimplex move and the reversible jump MCMC proposals. These are described in detail in the main text. MCMC analyses were run in RevBayes for 11000 iterations, where each iteration consisted of 79 MCMC moves per iteration. The 79 moves were randomly drawn from the 28 different Metropolis-Hastings moves listed in Table 1 using the weights listed. Samples of parameter values and joint ancestral states were drawn each iteration, and the first 1000 samples were discarded as burn in.

**Table 1:** MCMC moves used for chromosome number evolution analyses. See the main text for further explanations of the moves used. Samples were drawn from the MCMC each iteration, where each iteration consisted of 28 different moves in a random move schedule with 79 moves per iteration.

| | Parameter | $X$ | Move | Weight |
| --- | --- | --- | --- | --- |
| Anagenetic | Chromosome gain rate | $\gamma_a$ | Scale($\lambda = 1$) | 2 |
| | Chromosome gain rate | $\gamma_a$ | Reduce/Augment | 2 |
| | Chromosome loss rate | $\delta_a$ | Scale($\lambda = 1$) | 2 |
| | Chromosome loss rate | $\delta_a$ | Reduce/Augment | 2 |
| | Polyploidization rate | $\rho_a$ | Scale($\lambda = 1$) | 2 |
| | Polyploidization rate | $\rho_a$ | Reduce/Augment | 2 |
| | Demi-polyploidization rate | $\eta_a$ | Scale($\lambda = 1$) | 2 |
| | Demi-polyploidization rate | $\eta_a$ | Reduce/Augment | 2 |
| | Linear component of gain rate | $\gamma_m$ | Slide($\delta = 0.1$) | 1 |
| | Linear component of gain rate | $\gamma_m$ | Slide($\delta = 0.001$) | 1 |
| | Linear component of gain rate | $\gamma_m$ | Reduce/Augment | 2 |
| | Linear component of loss rate | $\delta_m$ | Slide($\delta = 0.1$) | 1 |
| | Linear component of loss rate | $\delta_m$ | Slide($\delta = 0.001$) | 1 |
| | Linear component of loss rate | $\delta_m$ | Reduce/Augment | 2 |
| Cladogenetic | No change | $\phi_c$ | Scale($\lambda = 5$) | 2 |
| | Chromosome gain | $\gamma_c$ | Scale($\lambda = 5$) | 2 |
| | Chromosome gain | $\gamma_c$ | Reduce/Augment | 2 |
| | Chromosome loss | $\delta_c$ | Scale($\lambda = 5$) | 2 |
| | Chromosome loss | $\delta_c$ | Reduce/Augment | 2 |
| | Polyploidization | $\rho_c$ | Scale($\lambda = 5$) | 2 |
| | Polyploidization | $\rho_c$ | Reduce/Augment | 2 |
| | Demi-polyploidization | $\eta_c$ | Scale($\lambda = 5$) | 2 |
| | Demi-polyploidization | $\eta_c$ | Reduce/Augment | 2 |
| | All cladogenetic rates | $\phi_c, \gamma_c, \delta_c, \rho_c, \eta_c$ | Joint Up-Down<br>Scale($\lambda = 0.5$) | 2 |
| Other | Root frequencies | $\pi$ | BetaSimplex($\alpha = 0.5$) | 10 |
| | Root frequencies | $\pi$ | ElementSwapSimplex | 20 |
| | Relative-extinction | $r$ | Scale($\lambda = 5$) | 3 |
| | Relative-extinction and all clado rates | $r, \phi_c, \gamma_c, \delta_c, \rho_c, \eta_c$ | Joint Up-Down<br>Scale($\lambda = 0.5$) | 2 |
| **Total** |  |  | **28** | **79** |

## Appendix 3: Simulation Details

### Description of Simulation Experiments

#### Experiment 1

In experiment 1 we tested the effect of unobserved speciation events due to extinction on chromosome number estimates when using a model that does not account for unobserved speciation. Is the additional model complexity required to account for unobserved speciation necessary, or are the effects of unobserved speciation negligible and safe to ignore? Using the non-SSE model described above that does not account for unobserved speciation, ancestral chromosome numbers and chromosome evolution model parameters were estimated for each of the 600 datasets.

#### Experiment 2

Here we compared the accuracy of models of chromosome evolution that account for unobserved speciation versus those that do not. Since extinction can safely be assumed to be present to some extent in all clades, it is likely that all empirical datasets contain some unobserved speciation. Do we see an increase in accuracy when we account for unobserved speciation events, or conversely do we see an increase in the variance of our estimates that perhaps describes true uncertainty due to extinction? To test this, we estimated ancestral chromosome numbers and chromosome evolution model parameters over the simulated datasets that included unobserved speciation using both ChromoSSE that accounts for unobserved speciation as well as the non-SSE model that does not.

#### Experiment 3

In experiment 3 we tested the effect of jointly estimating speciation and extinction rates with chromosome number evolution. Estimating speciation and extinction rates accurately is notoriously challenging (Nee et al. 1994; Rabosky 2010; Beaulieu and O’Meara 2015; May et al. 2016), so how much of the variance in chromosome evolution estimates made with models that jointly estimate speciation and extinction are due to uncertainty in diversification rates? Here we compared our estimates of ancestral chromosome numbers and chromosome evolution model parameters using ChromoSSE that accounts for unobserved speciation (and in which speciation and extinction rates are jointly estimated) with estimates made from ChromoSSE but where the true rates of speciation and extinction used to simulate the data were fixed. The latter analyses were given the true rates of total speciation and extinction, but still had to estimate the proportion of speciation events for each type of cladogenetic event.

#### Experiment 4

Since we model the same chromosome number transitions as both cladogenetic and anagenetic processes, it is possible that the two processes could be confounded and our models may not be fully identifiable. Furthermore, preliminary results suggested our models overestimate anagenetic changes and underestimate cladogenetic changes when the true generating process includes cladogenetic evolution. Here we compared cladogenetic and anagenetic estimates made by ChromoSSE under simulation scenarios that only included cladogenetic changes. Do we see an increase in accuracy of cladogenetic parameter estimates when anagenetic changes are disallowed (fixed to 0)?

**Table 2:** Simulation parameter values. Parameter values used to simulate datasets. The top 3 rows show the 3 modes of chromosome number evolution simulated for Experiments 1, 2, 3, and 4: anagenetic only, cladogenetic only, and mixed. Row 4 shows the parameter values used to simulate data for Experiment 5. The total speciation rate *λ_t_* = 0.25 and the extinction rate *μ* = 0.15. The root state was fixed to 8.

| Simulation mode | $\gamma_a$ | $\delta_a$ | $\rho_a$ | $\eta_a$ | $\gamma_m$ | $\delta_m$ | $\phi_c$ | $\gamma_c$ | $\delta_c$ | $\rho_c$ | $\eta_c$ |
| --- | --- | --- | --- | --- | --- | --- | --- | --- | --- | --- | --- |
| Anagenetic | 0.0085 | 0.0085 | 0.0085 | - | - | - | $\lambda_t$ | - | - | - | - |
| Cladogenetic | - | - | - | - | - | - | $0.85\lambda_t$ | $0.05\lambda_t$ | $0.05\lambda_t$ | $0.05\lambda_t$ | - |
| Mixed | 0.0085 | 0.0085 | 0.0085 | - | - | - | $0.85\lambda_t$ | $0.05\lambda_t$ | $0.05\lambda_t$ | $0.05\lambda_t$ | - |
| Experiment 5 | 0.0025 | 0.0025 | 0.0025 | - | - | - | $0.93\lambda_t$ | $0.02\lambda_t$ | $0.02\lambda_t$ | $0.02\lambda_t$ | - |

#### Experiment 5

Experiments 1-3 deal with the increase in uncertainty caused by unobserved speciation events due to extinction. Here we focused on the effect of unobserved speciation due to incomplete taxon sampling by comparing chromosome number estimates at 3 levels of taxon sampling: 100%, 50%, and 10%. We compared estimates made by both the ChromoSSE model and the non-SSE model, as well as compared estimates made by ChromoSSE using the true taxon sampling probability *ρ_s_* versus estimates made by ChromoSSE using *ρ_s_* fixed to 1.0.

### Methods Used to Simulate Data

For experiments 1, 2, 3, and 4 the same set of simulated trees and chromosome counts were used. Since ChromoSSE assumes the total rates of speciation and extinction are fixed over the tree (see Equation 5 of the main text), trees were first simulated with constant diversification rates, and then cladogenetic and anagenetic chromosome evolution was simulated over the trees. 100 trees were simulated under the birth-death process with *λ* = 0.25 and *μ* = 0.15 (see Figure 4) using the R package diversitree (FitzJohn 2012). The trees were conditioned on an age of 25.0 time units and a minimum of 10 extant lineages. To test the effect of unobserved speciation events due to lineages going extinct on cladogenetic estimates, chromosome number evolution was simulated along the trees including their extinct lineages (unpruned) and the same 100 trees but with the extinct lineages pruned. All chromosome number simulations were performed using RevBayes (Höhna et al. 2016).

**Figure 4:**
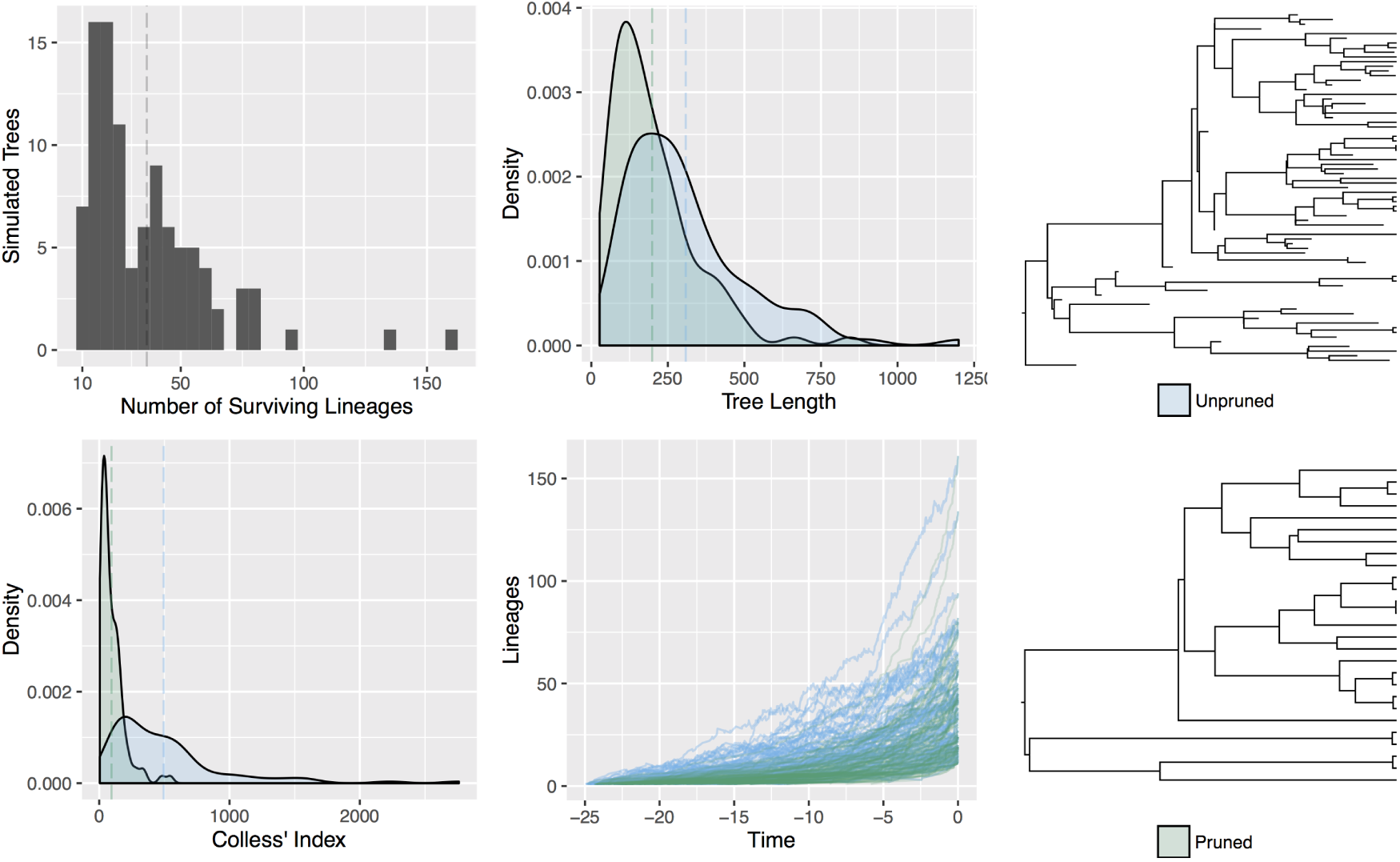
Tree simulations. 100 trees were simulated under the birth-death process as described in the main text for Experiments 1, 2, 3, and 4. Chromosome number evolution was simulated over the unpruned trees that included all extinct lineages, as well as over the same trees but with extinct lineages pruned. This resulted in two simulated datasets: one simulated under a process that did have unobserved speciation events, and one simulated with no unobserved speciation events. Shown above is a histogram of the number of lineages that survived to the present, the tree lengths, Colless’ Index (a measure of tree imbalance; Colless 1982), and lineage through time plots of the 100 pruned and unpruned trees.

Three models were used to generate simulated chromosome counts: a model where all chromosome evolution was anagenetic, a model where all chromosome evolution was cladogenetic, and a model that mixed both anagenetic and cladogenetic changes (Table 2). Parameter values were roughly informed by the mean values estimated from the empirical datasets. The mean length of the simulated trees was 253.5 (Figure 4). Hence, the anagenetic rates were set to 2/235.5 ≈ 0.0085 which corresponds to an expected value of 2 events over the tree for each of the four transition types. The root chromosome number was fixed to be 8. Simulating data for all 3 models over both the pruned and unpruned tree resulted in 600 simulated datasets. To reproduce the effect of using reconstructed phylogenies all inferences were performed using the trees with extinct lineages pruned and with chromosome counts from extinct lineages removed.

Since Experiment 5 focused on the effect of incomplete taxon sampling on chromosome number estimates, the trees used needed to be conditioned on a known number of extant tips. The trees used for the previous simulations were conditioned only on age and a minimum of 10 extant lineages and so were not appropriate. To simulate 100 trees conditioned on 200 extant lineages we used the R package TreeSim (Stadler 2011) with *λ* = 0.25 and *μ* = 0.15 (like above). Complete trees with both extant and extinct lineages were simulated, and then chromosome evolution was simulated over the complete tree. Since these trees had a significantly longer mean length (2020.1 compared to 253.5) we used different rates of chromosome evolution to simulate data compared to Experiments 1, 2, 3, and 4 (Table 2). Chromosome numbers were only simulated using a mixed anagenetic and cladogenetic model. The anagenetic rates were set to 5/2020.1 ≈ 0.0025 which corresponds to an expected value of 5 events over the tree for each of the four transition types. Like Experiments 1, 2, 3, and 4, the root chromosome number was fixed to be 8. Once chromosome data was simulated over the complete trees, the extinct taxa were pruned off leaving trees with 100% taxon sampling. 50% of the tips were randomly pruned off to create trees with 50% taxon sampling, and 90% of the tips were randomly pruned off to create trees with 10% taxon sampling.

## Appendix 4: MCMC Convergence of Simulation Replicates

Effective sample sizes (ESS) for all parameters in all simulation replicates were over 200, and the mean ESS values of the posterior for the replicates was 1470.3. Since the space of possible models is so large (1024 possible models, see main text), we replicated all analyses that included unobserved speciation in Experiment 1 three independent times to ensure that MCMC convergence was not an issue in detecting the true model of chromosome number evolution used to simulate the data. The results displayed in Table 3 show that the percentage of simulation replicates in which the true model was inferred to be the MAP model, and the mean posterior of the true model, converged and were stable across all three independent runs.

**Table 3:** Simulation Experiment 1 replicated 3 times. Estimates of the true model that generated the simulated data and estimates of the posterior probability of the true model were stable and converged across multiple independent replicates of the experiment.

| Replicate | Mode of Evolution Used to Simulate Data | True Model Estimated (%) | Mean Posterior of True Model |
| --- | --- | --- | --- |
| 1 | Cladogenetic | 15 | 0.09 |
| 1 | Anagenetic | 36 | 0.12 |
| 1 | Mixed | 2 | 0.10 |
| 2 | Cladogenetic | 15 | 0.09 |
| 2 | Anagenetic | 36 | 0.12 |
| 2 | Mixed | 2 | 0.09 |
| 3 | Cladogenetic | 15 | 0.09 |
| 3 | Anagenetic | 36 | 0.12 |
| 3 | Mixed | 2 | 0.10 |

